# Long-lived chondroprogenitors are generated by Gli1^+^ fetal-limb cells and are replenished upon cell-cycle arrest in the cartilage

**DOI:** 10.1101/2024.07.29.603524

**Authors:** Xinli Qu, Ehsan Razmara, Chee Ho H’ng, Kailash K. Vinu, Luciano G. Martelotto, Magnus Zethoven, Fernando J. Rossello, Shanika L. Amarasinghe, David R. Powell, Alberto Rosello-Diez

**Affiliations:** Australian Regenerative Medicine Institute, Monash University, Clayton, Victoria, Australia; University of Melbourne Centre For Cancer Research, The University of Melbourne, Melbourne, Victoria, Australia; Adelaide Centre for Epigenetics. South Australian immunoGENomics Cancer Institute. University of Adelaide, Adelaide, South Australia, Australia; Peter MacCallum Cancer Centre, Melbourne, Victoria, Australia; Murdoch Children’s Research Institute, The Royal Children’s Hospital, Melbourne, VIC 3052, Australia; Novo Nordisk Foundation Center for Stem Cell Medicine, Murdoch Children’s Research Institute, Melbourne, VIC 3052, Australia; Department of Clinical Pathology, University of Melbourne, Melbourne, VIC, Australia; Monash Genomics and Bioinformatics Platform, Monash University, Clayton, Victoria, Australia.; Present address: Biomedicine Discovery Institute, Monash University, Clayton, Victoria, Australia.; Monash eResearch Centre, Monash University, Clayton, Victoria, Australia; Department of Physiology, Development and Neuroscience, University of Cambridge, Cambridge, UK; Department of Genetics, University of Cambridge, Cambridge, UK; Centre for Trophoblast Research, University of Cambridge, Cambridge, UK

**Keywords:** Gli1, Pdgfra, growth plate, endochondral ossification, chondroprogenitors, long bones, limb growth, skeletal stem cells, perichondrium, catch-up growth, developmental robustness

## Abstract

The growth-plate cartilage of the developing long bones is a well-known system of spatially segregated stem/progenitor, transient amplifying and terminally differentiated cells. However, the regulation of the number and activity of long-lived cartilage progenitors (LLCPs) is poorly understood, despite its relevance for understanding human-height variation, the evolution of limb size and proportions and the aetiology of skeletal growth disorders. Moreover, whether their behaviour can adapt to developmental perturbations, generating robustness, has not been explored. Here, we show that Gli1^+^ cells are the fetal precursors of postnatal LLCPs. Moreover, while during normal growth they remain mostly dormant until postnatal stages, in response to cartilage-targeted cell-cycle arrest they increase their cartilage contribution, compensating for the perturbation. We further show that, during the response to challenge, some of the reparative Gli1^+^ cells originate from Pdgfra^+^ cells outside the cartilage anlage. Elucidating how stromal cells become Gli1^+^ LLCPs could shed light on developmental robustness and lead to growth-boosting therapies.

## Introduction

Developmental systems often display remarkable robustness, that is, the ability to overcome challenges that affect cell function or viability^1^. Elucidating the mechanisms underlying robustness could improve our fundamental understanding of how individual cells integrate information and coordinate with their neighbours such that organ-level collective behaviours are achieved. Moreover, it could lead to growth- modulating therapies for congenital or acquired growth disorders. Long bones in the limbs are a powerful model to study growth regulation. They grow via a transient cartilage template that is progressively replaced by bone^2^. Cartilage cells (chondrocytes) undergo progressive end-to-centre differentiation, starting as resting/reserve chondrocytes that are progressively recruited into the proliferative pool (flat cells arrayed in columns), which eventually enlarge and differentiate to hypertrophic chondrocytes. Some hypertrophic chondrocytes die, while others transdifferentiate into osteoblasts (bone-laying cells) that replace the cartilage with bone, forming the primary ossification center^3,4^. Osteoblasts can also migrate, along with blood vessels, from the perichondrium^5^, a connective tissue layer that surrounds the cartilage and has been shown to contain a stem-cell niche known as groove of Ranvier^6^. A secondary ossification centre (SOC) forms later (∼postnatal day 7 in the mouse) at both ends of the skeletal element, so that the growth cartilage gets ‘sandwiched’ into a so-called growth plate by both ossification centres.

Maintaining the size and structure of the growth plate–in other words, the balance between cartilage generation and destruction–is critical to sustain bone growth over time and requires refined control mechanisms. The size of the proliferative zone is maintained approximately constant due to a well- described negative feedback loop between Indian hedgehog and Parathyroid hormone related peptide that balances proliferation and differentiation^3^. But other control mechanisms are less known, especially regarding the resting-to-proliferative transition. In fetuses and neonates, cartilage progenitors (CPs) located in the reserve zone do not self renew^7^, so that when one is recruited into a proliferative column, its lineage gets rapidly exhausted, leading to short clonal columns. At approximately three weeks of age, CPs start to self-renew–possibly in response to SOC signals–giving rise to long clonal columns (Ext. Data Fig. 1)^7^. These long-lived CPs (LLCPs) are the key driver of subsequent growth and therefore of final bone length. Therefore, elucidating their origin and regulation is critical to understanding normal and compensatory growth, as well as skeletal growth disorders and the evolution of limb size. One of the biggest unresolved questions is whether postnatal LLCPs arise from fetal short-lived ones, in response to intrinsic and/or extrinsic triggers (Ext. Data Fig. 1a), or whether they are already present but inactive in the fetal limb, becoming activated (or even recruited from outside the cartilage) in response to cartilage maturation (Ext. Data Fig. 1b). Determining which of these mechanisms take place during normal growth, and whether they can adapt in response to developmental perturbations, is desirable for the development of targeted growth therapies, focussed on stimulating the expansion of the pool of LLCPs at the right time and location. In this study, we probed the system using models of cartilage- targeted cell-cycle arrest, combined with lineage tracing and functional analyses. We found that Gli1^+^ cells in the fetal limb give rise to dormant CPs, which later postnatally give rise to most of the chondrocytes of the growth plate. Moreover, these progenitors can be precociously activated by cartilage-targeted growth challenges, revealing significant plasticity and a strong reparative potential. We posit that this fundamental knowledge will be critical to understanding and manipulating cartilage growth and repair.

**Figure 1.**
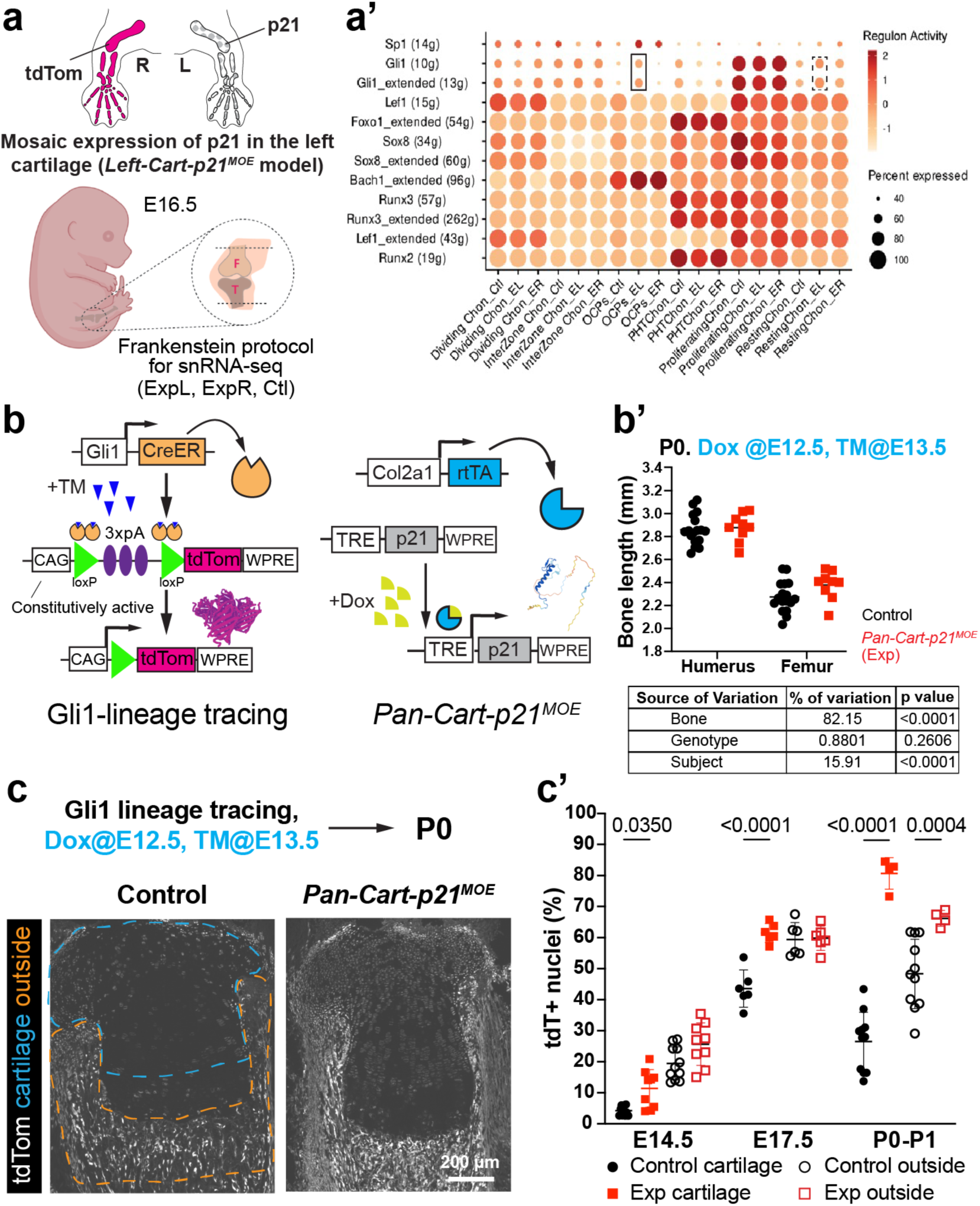
Mosaic cell-cycle arrest in the fetal cartilage triggers compensation from Gli1^+^ stromal cells. **a**, **a’**, A left-specific, cartilage-targeted p21 mosaic over-expression model (*Left-Cart-p21^MOE^*) was used for snRNA-seq analysis of knees. Left (L) and right (R) knees were processed separately but analysed together in control embryos (a). Chondrocyte-related cells were analysed via SCENIC and relevant regulons are shown boxed (a’, dashed lines denote mild changes). **b**, **b’**, Expression of p21 in the cartilage, induced from E12.5 to P0 (b), does not lead to growth defects (b’). The table shows results of 2-way ANOVA. **c**, **c’**, The Gli1 lineage was followed in absence (Control) and presence of p21 misexpression (*Pan-Cart-p21^MOE^*), and quantified in the regions shown in (c). In c’, data shown as mean± SD. p-values for Sidak’s multiple comparisons test after 2-way ANOVA are shown.

## Results

### Mosaic cell-cycle arrest in the fetal cartilage triggers a compensatory response from Gli1^+^ cells

We previously generated a fetal mouse model of <u>left</u>-<u>cart</u>ilage targeted, <u>m</u>osaic <u>o</u>ver<u>e</u>xpression of <u>p21</u> (which blocks the cell cycle in G1 phase^8^), by combining *Pitx2-Cre*^9^, *Col2a1-rtTA*^10^ and *Tigre^Dragon-p^*^21^ alleles^11^ (Ext. Data Fig. 2a, *Left-Cart-p21^MOE^* model hereafter). In that study, we showed that the mosaic cell-cycle arrest was compensated by hyperproliferation of the spared (p21^−^) chondrocytes^11^. We reasoned that, if short-lived CPs were the only source of LLCPs (as in Ext. Data Fig. 1a), their hastened proliferation would lead to premature exhaustion of the CP pool (Ext. Data Fig. 2b), and therefore to reduced bone length in the long term. To test this hypothesis, we followed mice from the *Left-Cart- p21^MOE^* model until postnatal day (P) 100, well beyond the end of the growth period. Interestingly, no major asymmetries in bone length were found as compared to *Pitx2-Cre/+; Tigre^Dragon-p^*^21^*^/+^* mice (Controls, Ctl hereafter, Ext. Data Fig. 2c, the left/right ratio was not significantly different for the tibia, and only ∼1.5% lower for the femur). This result suggested that either the spared resident CPs in the *Left-Cart-p21^MOE^*acquired self-renewing properties precociously, or that facultative CPs (i.e. cells that would not normally contribute to cartilage at that age) were recruited to the challenged cartilage. To shed light into this process, we undertook an unbiased single-nuclei RNA-seq approach of left and right fetal knees from two E16.5 Ctl and two *Left-Cart-p21^MOE^* embryos (Fig. 1a and Ext. Data Fig. 3a). Out of the original dataset (∼32,000 nuclei in 32 clusters, Ext. Data Fig. 3b, d), which contained mostly lateral plate mesoderm-derived cells, and a minority of muscle and endothelial cells, we further explored the former population (∼21,000 nuclei, Ext. Data Fig. 3d’). This subset was composed of 23 clusters (Ext. Data Fig. 3c, d), which we grouped in five main categories (see Methods): chondrocytes (Ext. Data Fig. 3b’), tenocytes, fibroblasts, perichondrium and joint-associated mesenchymal cells. Analysis of regulon activity with SCENIC^12^ revealed that the activity of GLI-Kruppel family member 1 (GLI1) transcription factor was significantly upregulated in the left experimental (ExpL) osteochondroprogenitors and to a lesser extent in resting chondrocytes (Fig. 1a’). This was interesting because Gli1^+^ osteochondroprogenitors have been shown to participate in the repair of bone fractures^13^. Thus, we next asked whether the number of Gli1^+^ cells changed in response to cartilage-targeted p21 overexpression. Indeed, the Propeller tool^14^ revealed that, within the Gli1^high^ lateral-plate-mesoderm-derived population, the proportion of cells belonging to the chondrocyte group was increased in experimental limbs, as compared to Ctl (Ext. Data Fig. 3e,e’). To confirm these findings, we turned to Cre-based lineage tracing. To trace the Gli1-lineage independently of p21 expression, we utilised a Doxycycline (Dox)-controlled, Cre-independent p21 expression system, targeted to all cartilage elements (*Col2a1-rtTA/+; Tigre^TRE-p^*^21^*^/+^*, *Pan-Cart-p21^MOE^* model hereafter; TRE, Tet-responsive element)^15^. Females bearing a tamoxifen (TM)-inducible Cre recombinase (CreER^T2^) cassette^16^ targeted to the *Gli1* locus (*Gli1^CreER^* hereafter) were crossed with *Pan-Cart-p21^MOE^*males bearing also a Cre-reporter cassette (lox-STOP-lox-tdTomato) targeted to the safe harbour locus *Rosa26* (*R26^LSL-tdTom^* hereafter, Fig. 1b)^17^ and given Dox at embryonic day (E) 12.5 and TM at E13.5. As with the model targeted to the left limb, we confirmed that expression of p21 in 60-70% of chondrocytes (Ext. Data Fig. 4a) did not cause growth defects by postnatal day (P) 0 (Fig. 1b’). Despite continuous Dox exposure, the fraction of p21^+^ chondrocytes decreased with time (Ext. Data Fig. 4a), as we previously described with the *Left-Cart-p21^MOE^* model^11^. To assess the involvement of Gli1^+^ cells in this compensatory response, we followed the distribution of tdTom^+^ cells inside and outside the cartilage 1, 4, 7 days post TM (dpt, Fig. 1c,c’). In *Pan-Cart-p21^MOE^*mice, the Gli1-lineage showed a significant cartilage expansion, as compared to controls, at all stages analysed (Fig. 1c’). These results suggested that, in control conditions, most E13.5-marked Gli1-derived chondrocytes only contribute transiently and are then “pushed out” of the cartilage, while in *Pan-Cart- p21^MOE^* limbs, the Gli1-lineage increases its contribution to longer-lived CPs. Given that CPs reside in the resting zone^18^, we divided the cartilage analysis into resting and proliferative, finding that both zones showed increased number of tdTom^+^ cells, typically biased towards the resting zone (Ext. Data Fig. 4b- b’’). Moreover, the contribution outside the cartilage was also expanded in P0 *Pan-Cart-p21^MOE^* limbs compared to controls (Fig. 1c’), suggesting that the challenged cartilage signals to the surrounding tissues.

**Figure 2.**
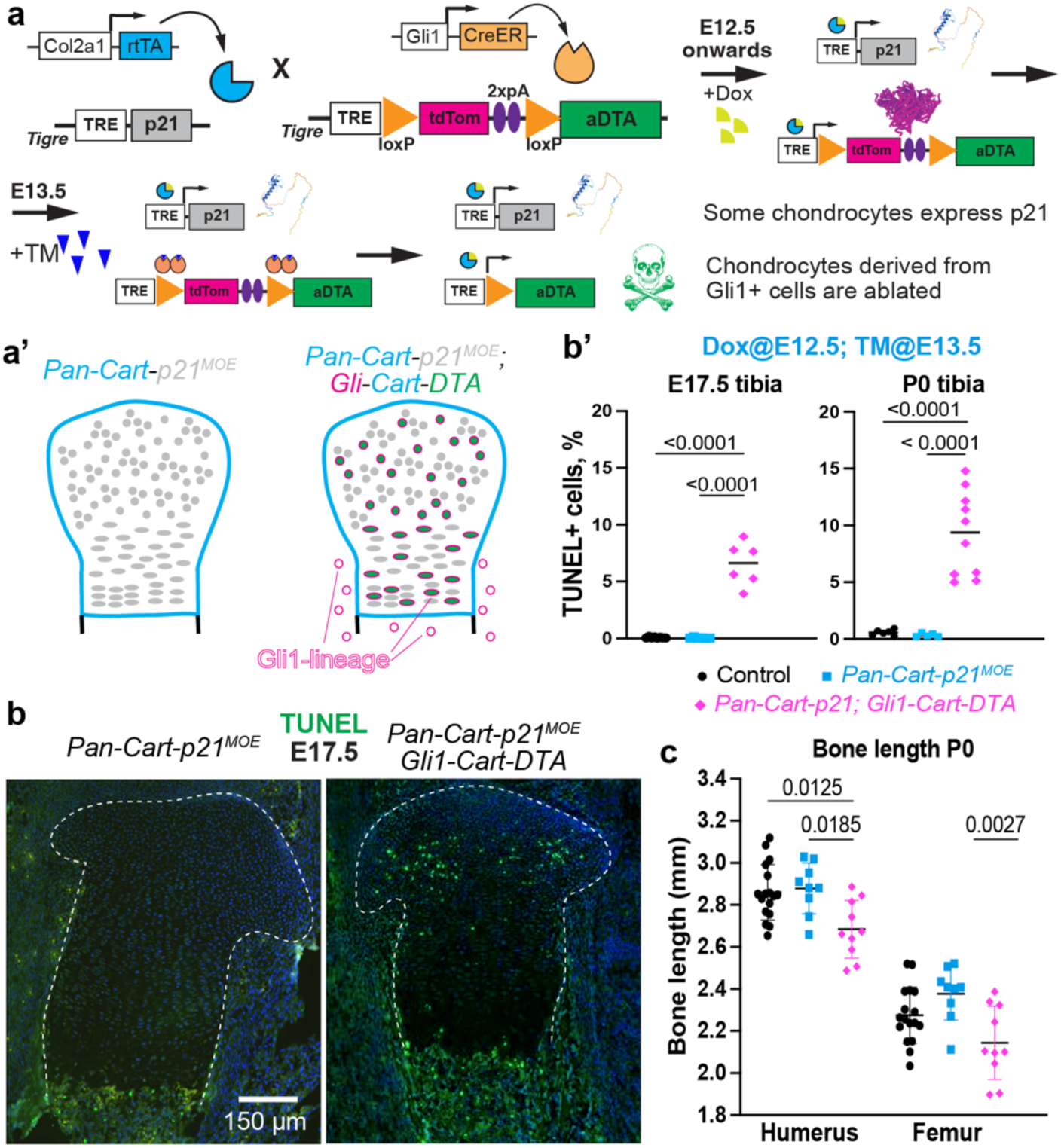
Gli1-derived chondrocytes are needed for compensation of *Pan-Cart-p21^MOE^*. **a**, A *Dragon-aDTA* allele (aDTA= attenuated diphtheria toxin) is combined with *Gli1^CreER^* and *Pan-Cart-p21^MOE^* to ablate Gli1-derived chondrocytes during p21-induced compensation. **b, b’**, Example images of TUNEL (b, cartilage outlined) and quantification (b’) for the indicated stages and genotypes (n=8,6,6 at E17.5, n=6,5,10 at P0). Significant p-values for Tukey’s test after ANOVA are shown. **c**, Length of the P0 mineralised region is shown for the indicated genotypes as mean±SD (n=17, 9, 10). Significant p-values of Sidak’s multiple comparisons test (after 2-way ANOVA) are shown.

**Figure 3.**
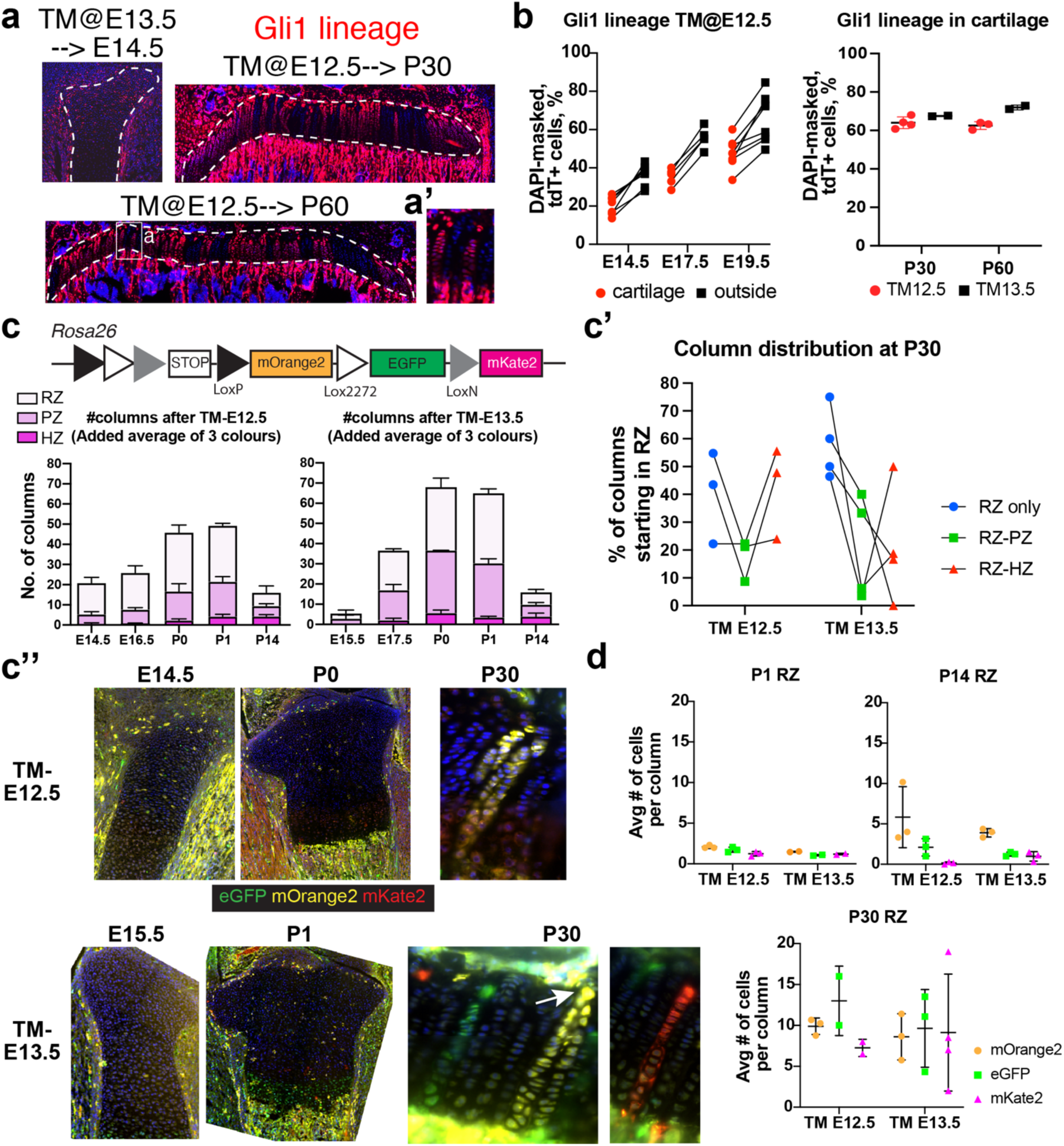
Fetal Gli1^+^ cells give rise to most chondrocytes in the juvenile and adult growth plate. **a**, Examples of Gli1-lineage tracing after TM induction at E12.5 or E13.5, analysed at the indicated stages. **b**, Quantification of reporter-expressing cells, inside and outside the cartilage for TM-E12.5 (left, n=6,5,6), or in cartilage for both inductions (right, n=4,2). **c**-**c’’**, The tri-colour reporter RGBow was used to quantify individual clones in the growth plate (c, added average of each colour is shown), at multiple times post induction (n=3, 4, 3, 3, 3, 3 for TM-E12.5, n=3, 3, 2, 2, 2, 3, 3 for TM-E13.5). RZ, PZ, HZ: resting, proliferative, hypertrophic zone. In c’, the % of P30 columns starting from RZ are shown, based on where they end (n= 3 and 4). c’’ shows representative images. Arrow: RZ-rooted LLCP. **d**, number of cells per RZ-rooted column, for the stages and induction times indicated.

**Figure 4.**
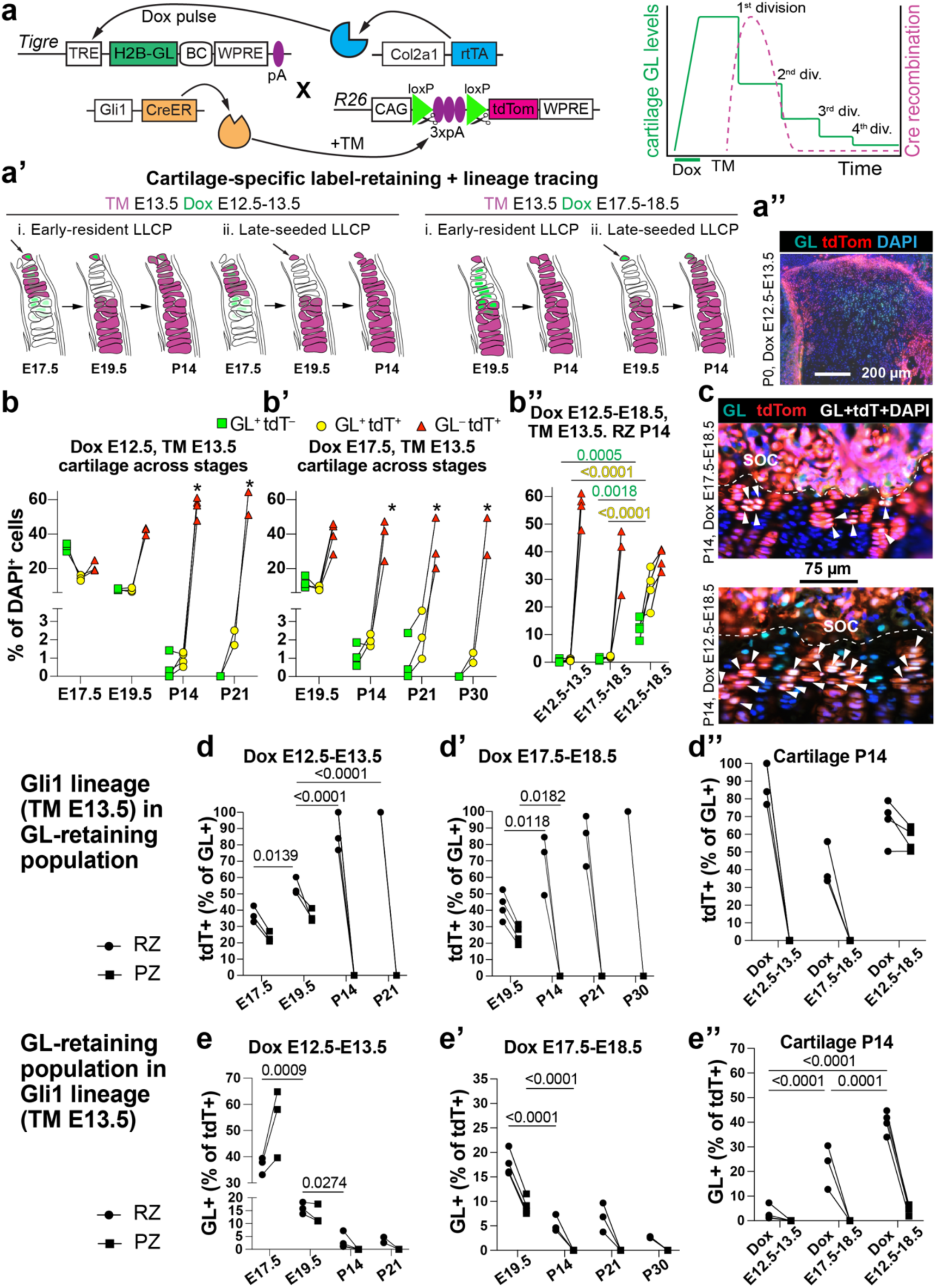
Most fetal LLCPs belong to the Gli1 lineage. **a**, **a’**, Model to permanently label the Gli1 lineage and transiently label chondrocytes with a nuclear GreenLantern (H2B-GL) that gets halved every cell division (a); different outcomes depending on Dox timing (a’) and a representative image of the distribution after early labelling (a’’). BC, barcoding region (not used here). **b**-**b’’**, Quantification of single-positive and double-positive populations in the cartilage at the indicated stages after induction at E12.5-E13.5 (b, n=3, 3, 4, 2), E17.5-E18.5 (b’, n=4, 3, 3, 2) or E12.5-E18.5 (b’’, n=4). *: quantified in resting zone only. **c**, Representative images of tdT and nGL overlap (arrowheads) at P14 with 2 different induction regimes. SOC, secondary ossification centre (separated from cartilage by dashed lines). **d**-**e’’**, % of tdT^+^ cells within the GL^+^ population (d-d’’) or GL^+^ cells within the tdT^+^ population (e-e’’) after 1-day Dox pulse at E12.5 (d, e) or E17.5 (d’, e’) or 6-day pulse at E12.5 (d’’, e’’). In all graphs p-values for multiple comparisons tests after 2-way ANOVA are shown.

Importantly, since our initial labelling of the Gli1-lineage (E13.5-TM) was done after the induction of p21 expression (E12.5), we were not able to distinguish between two scenarios: i) all the Gli1^+^ cells derive from formerly Gli1^+^ cells; ii) *Gli1* expression is activated *de novo* in cells that were Gli1^−^ before. Since the second scenario would reveal an unknown degree of plasticity that could be relevant for the understanding of growth control and the design of future therapies, we decided to further investigate this matter. We reasoned that, by providing Dox and TM both at E12.5, most of the tdTom-labelling would happen among existing Gli1^+^ cells, with most of the TM being washed out by the time any potential *de novo* expression of Gli1 took place. Using this approach, we found that the number of descendants was lower in P0 *Pan-Cart-p21^MOE^*as compared to E12.5-Dox/E13.5-TM (Ext. Data Fig. 4c-c’). This result suggests that while the existing population of Gli1^+^ cells contributes to the repair, some Gli1^−^ cells become Gli1^+^ <u>after</u> *p21* induction in the cartilage.

To test whether Gli1-derived chondrocytes are required for the compensation of *Pan-Cart-p21^MOE^*, we used a *Tigre^Dragon-aDTA^*allele in which inducible expression of aDTA (attenuated diphtheria toxin) requires Cre and rtTA activity^15,19^ (Fig. 2a). Mice were crossed as shown in Fig. 2a to generate *Gli1^CreER/+^*; *Col2a1-rtTA/+; Tigre^TRE-p^*^21^*^/+^* embryos (phenotypically akin to *Pan-Cart-p21^MOE^*) and *Gli1^CreER/+^; Col2a1-rtTA/+; Tigre^Dragon-aDTA/TRE-p^*^21^ littermates (*Pan-Cart-p21^MOE^; Gli-Cart-DTA,* Fig. 2a), in addition to control (rtTA^−^) ones. We gave Dox from E12.5 till sample collection to induce p21 expression in the cartilage (*Pan-Cart-p21^MOE^*), and TM at E13.5 to induce cell death of the rtTA- expressing Gli1-derived chondrocytes (*Gli-Cart-DTA* combination). As predicted, TUNEL staining revealed apoptosis only in the cartilage of *Pan-Cart-p21^MOE^; Gli-Cart-DTA* embryos (Fig. 2b, b’). We then measured bone length at P0 and found that bone growth was impaired in *Pan-Cart-p21^MOE^; Gli- Cart-DTA* pups as compared to *Pan-Cart-p21^MOE^* and/or control pups (Fig. 2c). This result demonstrates that Gli1-derived chondrocytes are required in the E13.5 to P0 period to compensate for cell-cycle arrest in the cartilage. Of note, due to increased mortality of *Pan-Cart-p21^MOE^; Gli1-Cart-DTA* mice, later time points could not be analysed.

### Fetal Gli1^+^ cells give rise to most chondrocytes in the juvenile and adult growth plate

Having identified a highly-reparative fetal Gli1^+^ population, we set out to determine their role during normal growth. When Gli1^+^ cells are lineage-traced at one month of age, they label 60-80% of chondrocytes of the growth plate one month later^20^, indicating that P30 Gli1^+^ chondrocytes contain long- lived chondroprogenitors (LLCPs). Since Gli1^+^ cells exist in the limb at fetal stages (Ext. Data Fig. 5a, a’), we tested whether fetal Gli1^+^ cells already contained LLCPs. We crossed *Gli1^CreER/CreER^*females with *R26^LSL-tdTom/LSL-tdTom^* males and provided TM at either E12.5 or E13.5 and quantified the number of tdTom^+^ cells at multiple stages (E14.5, E17.5, E19.5 and P30 and P60). In all cases, tdTom^+^ cells were found both inside the cartilage and adjacent tissues, including perichondrium, metaphysis, interzone knee region and SOC (Fig. 3a, a’). Since fetal CPs had been shown not to self-renew^7^, we expected the % of labelled chondrocytes to decrease with time. On the contrary, the contribution of Gli1-lineage cells to the cartilage increased with time, labelling up to 65-70% of chondrocytes in the 2-month-old growth plate (Fig. 3b). To ascertain when in this process Gli1-derived chondrocytes switched from forming short to forming long clonal columns, we used a tri-colour reporter line (RGBow, Fig. 3c)^21^ that randomly expresses one of three membrane-bound fluorescent proteins upon Cre-recombination. After giving TM at E12.5 or E13.5, we analysed the number, location and length of distinct columns in the three regions of the cartilage. Notably, the total number of columns increased up to P0-P1, the majority located in the resting and proliferative zones (Fig. 3c, c’’) and composed of 2-3 cells (Fig. 3d). By P14, the total number of columns was significantly reduced but started to become longer (Fig. 3c’’, d). By P30 few columns remained, but many crossed the whole growth plate starting from the RZ (Fig. 3c’, c’’). In summary, the results indicated that Gli1^+^ cells at E12.5 or E13.5 give rise to CPs with two different fates: short-lived clones that get depleted over time, and long-lived ones that remain dormant at peri-natal stages and become more active after the second postnatal week.

**Figure 5.**
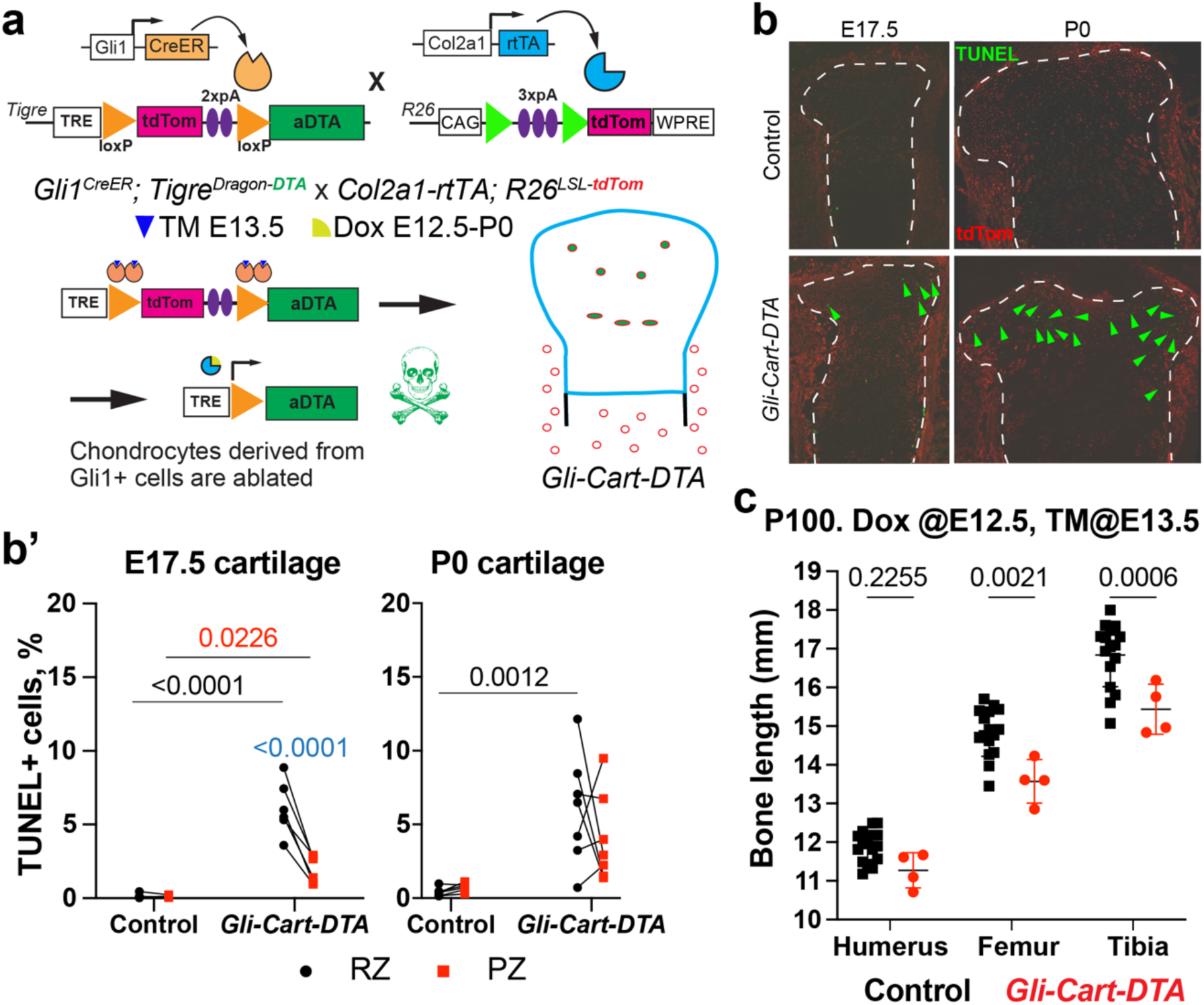
**a**, The *Dragon-aDTA* allele was combined with *Gli1^CreER^* and *Col2a1-rtTA* to activate diphtheria toxin expression exclusively in Gli1-lineage chondrocytes in the last week of gestation (*Gli-Cart-DTA* model). **b**, **b’** Typical images (b) and quantification (b‘) of TUNEL^+^ cells (arrowheads) at the indicated stages. **c** Bone length of Ctl and *Gli-Cart-DTA* mice at postnatal day 100 (P100). In b’ and c, p-values of multiple comparisons tests after 2-way ANOVA are shown. RZ, PZ: resting proliferative zone.

### Long-lived CPs are derived from Gli1^+^ cells that continuously seed the fetal and perinatal cartilage

We next asked whether Gli1-derived LLCPs were already present in the early cartilage template (around E12.5), or whether they were incorporated later from the adjacent tissues. The latter possibility is supported by studies showing that the chicken perichondrium is a reservoir for cartilage precursors^22^. To this end, we generated a mouse line in which expression of nuclear-directed Green Lantern fluorescent protein depends on Dox-activated rtTA (Fig. 4a and Ext. Data Fig. 6a-b, *Tigre^TRE-nGL^* hereafter). We then crossed *Gli1^CreER/CreER^; Tigre^TRE-nGL/TRE-nGL^* females with *Col2a1-rtTA/+; R26^LSL-tdTom/LSL-tdTom^* males, and provided TM at E13.5 to label the Gli1 lineage with tdTom, and a 1-day pulse of Dox to transiently express nGL primarily in fetal chondrocytes. We then followed the green signal over time, with the expectation that quiescent chondrocytes (that is, the cells that will become LLCPs) would retain fluorescence for longer than cells dividing quickly (Fig. 4a). Moreover, by providing Dox at different points, we aimed to label different potential CP populations (Fig. 4a’): those already resident in the early cartilage anlage (Dox at E12.5), and those seeded later into the cartilage (Dox at E17.5). Most green cells after E12.5-Dox were found inside the cartilage, with a negligible number in the perichondrium (Fig. 4a’’, Ext. Data Fig. 6c-c’), confirming that the new tool could be used to track cells located only in the cartilage. As predicted, the number of GL^+^ chondrocytes dropped fast at first (∼2.5-fold from E17.5 to E19.5), but then stabilised at 2-3% of the RZ cells at P14 and P21 (Fig. 4b, no green signal in the PZ), becoming undetectable by P30. Remarkably, the % of Gli1-lineage chondrocytes in the GL^+^ population increased over time in the RZ (Fig. 4d), until ∼90% of GL-retaining CPs at P14 belonged to the E13.5-labelled Gli1-lineage. Conversely, only ∼3% of the Gli1-lineage cells in the P14 RZ were also GL^+^, indicating that the Gli1-lineage contains other subpopulations (Fig. 4e). Similar results were obtained when Dox was given between E17.5 and E18.5 (e.g. Fig. 4b’, c, d’, e’), except for the proportion of GL^+^ cells within the Gli1-lineage RZ chondrocytes (∼6.7% at P21). These results suggested that while many LLCPs are already present in the E12.5 cartilage, more LLCPs are added later, in both cases derived mostly from fetal Gli1^+^ cells. To test this prediction, we extended the Dox pulse from E12.5 to E18.5, so that newly added chondrocyte waves were also labelled. As predicted, many more double positive cells were found in the P14 RZ than with any other regimen (Fig. 4b’’, c, d’’, e’’), suggesting that from E12.5 to E18.5 there is a continuous addition of LLCP precursors from outside the cartilage.

**Figure 6.**
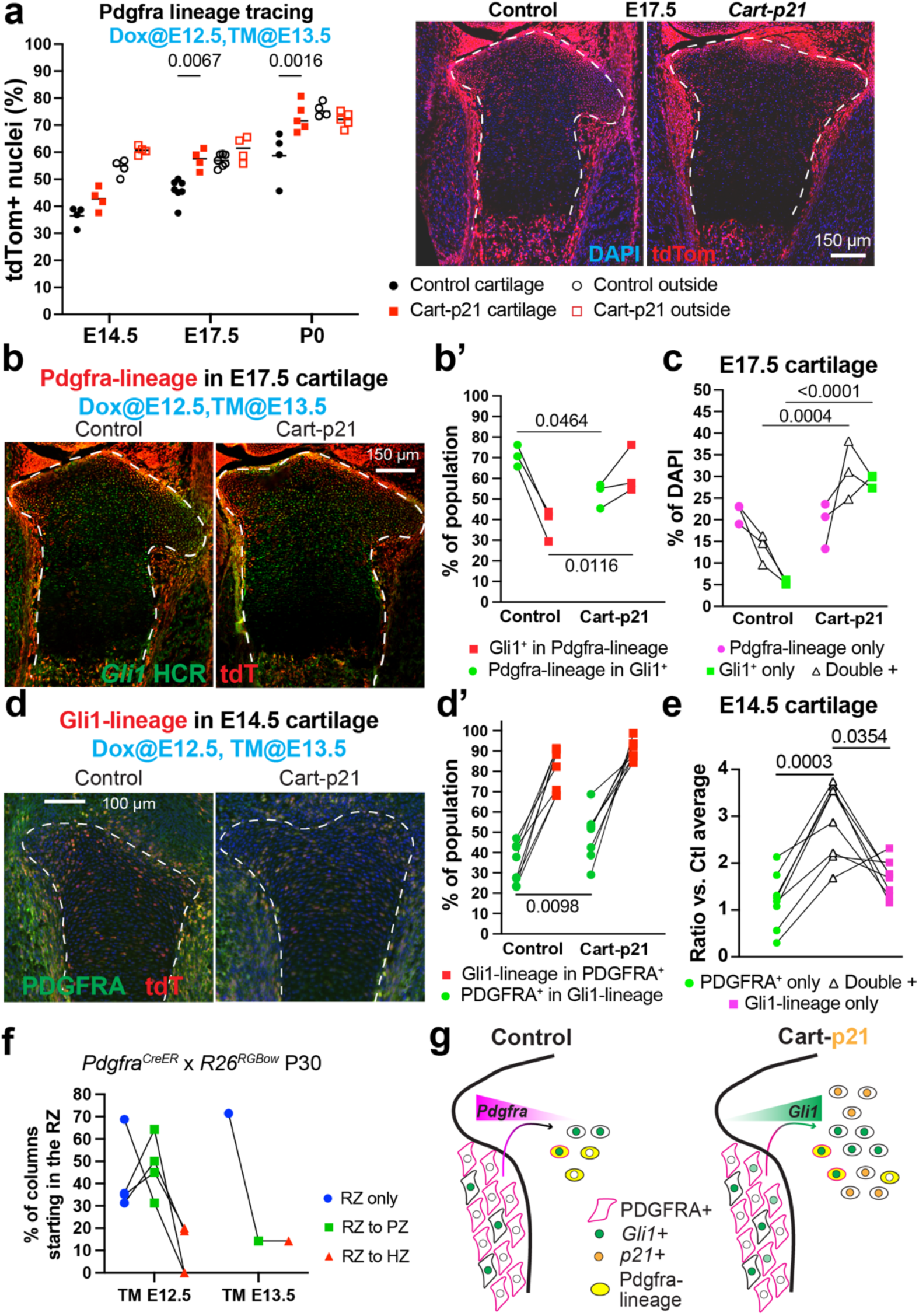
The contribution of the Pdgfra lineage to Gli1^+^ LLCPs increases upon cartilage challenge. **a**, The Pdgfra lineage was traced in the absence and presence of *Pan-Cart-p21^MOE^*, both inside and outside the cartilage (dashed lines; n=4 Ctl, 4 Exp at E14.5; 6, 4 at E17.5; 4, 5 at P0). **b**-**c**, Representative images (b) and quantification (b’, c) of Pdgfra-lineage^+^ and/or Gli1^+^ cells in the E17.5 cartilage (n=3 Ctl, 3 Exp). **d**-**d’**, Representative images (d) and quantification (d’) of Gli1-lineage^+^ and/or PDGFRA^+^ cells in the E14.5 cartilage (n=5 Ctl, 6 Exp). **e**, Ratio of the number of PDGFRA^+^ and/or Gli1-lineage^+^ cells in experimental cartilage, divided over the average of the same measurement in control samples. **f**, RGBow-expression was induced in Pdgfra^+^ cells at either E12.5 (n=4) or E13.5 (n=1) and followed till P30. RZ, PZ, HZ: resting, proliferative, hypertrophic zone. The % of columns starting from RZ are shown, based on where they end. In a, b’, c, d’, e, p-values for Sidak’s posthoc test after 2-way ANOVA. **g**, Potential model of how Pdgfra-lineage moving into the cartilage could become Gli1^+^ LLCPs in response to *Pan-Cart-p21^MOE^*.

### Chondrocytes generated in the last postnatal week from Gli1+ cells are required for long-term bone growth

To test if Gli1-derived chondrocytes are required to sustain bone growth, we crossed *Gli1^CreER^; Tigre ^Dragon-aDTA^* mice with *Col2a1-rtTA; R26^LSL-tdTom^*, and gave TM at E13.5 and Dox E12.5-P0. This way, only chondrocytes were ablated among Gli1-descendants (Fig. 5a-b’). While the ablation was incomplete and only sustained for a short time, this targeted injury led to reduced bone length by P100, especially for the tibia (Fig. 5c). This revealed the important role of fetal/perinatal Gli1-derived chondrocytes in long-term bone growth. Of note, ablation could not be sustained postnatally due to insufficient activity of *Col2a1-rtTA*, raising the possibility that partial compensation had taken place postnatally.

### Some of the Gli1^+^ LLCPs generated upon challenge derive from stromal Pdgfra^+^ cells

Having found evidence suggesting that stromal (non-cartilage) Gli1^+^ cells can become LLCPs (Fig. 4), and that some Gli1^−^ cells become Gli1^+^ upon p21 misexpression (Ext. Data Fig. 4), we next set out to determine the origin and regulation of Gli1^+^ cells in normal and challenged growth.

To capture the potential arising of new Gli1^+^ chondrocytes from stromal cells, we chose *platelet derived growth factor alpha* (*Pdgfra*) as a marker of stromal cells that is not expressed in the cartilage (Ext. Data Fig. 5b, b’). We crossed *Pdgfra^CreER/+^*females^23^ with *Pan-Cart-p21^MOE^*; *R26^LSL-tdT/LSL-tdT^*males and provided Dox at E12.5 and TM at E13.5. We found that, in normal growth, the E13.5-labelled Pdgfra- lineage gave rise to an increasing proportion of chondrocytes (Fig. 6a, 30-40% at E14.5, 45-50% at E17.5 and ∼60% at P0), strongly suggesting that the surrounding tissues can contribute to cartilage. Moreover, the proportion of Pdgfra-lineage chondrocytes was progressively increased in *Pan-Cart- p21^MOE^*limbs as compared to controls (Fig. 6a, compare P0 with E14.5). Of note, the Pdgfra-lineage located in the resting zone showed increased proliferation in *Pan-Cart-p21^MOE^* vs. control limbs at E14.5 and E17.5 (Ext. Data Fig. 7a-b’), potentially explaining the progressive expansion observed at and after E17.5 (Fig. 6a).

**Figure 7.**
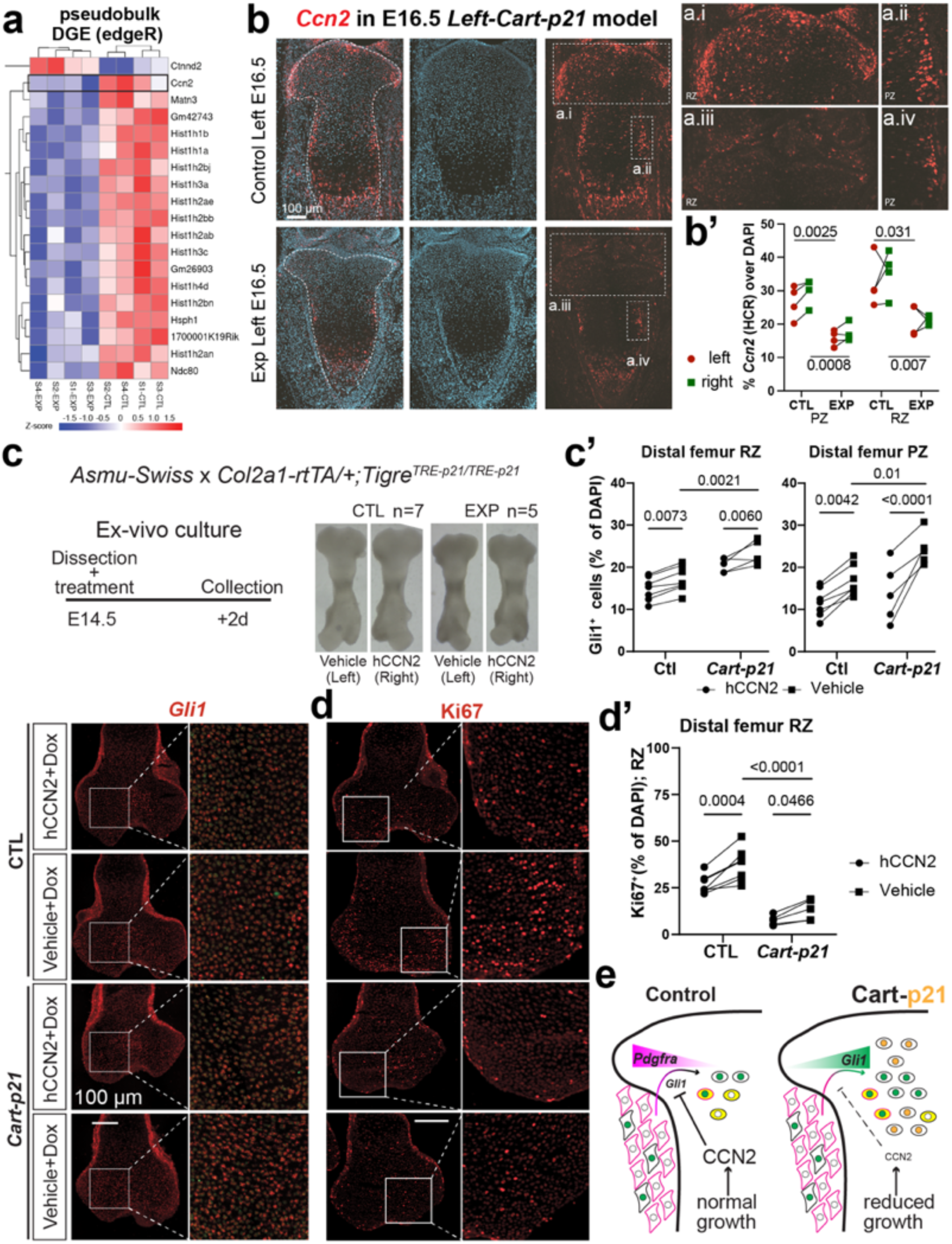
CCN2 impairs *Gli1* expression and proliferation in chondrocytes, and is downregulated in *Cart-p21^MOE^* samples. **a**, <u>D</u>ifferential <u>G</u>ene <u>E</u>xpression analysis (pseudobulk) of scRNA-seq data using edgeR. **b**, **b’**, *Ccn2* expression (HCR) in control and *Left-Cart-p21^MOE^* E16.5 samples (a), as indicated, and quantified in (b). n=4 CTL and 4 EXP embryos. Boxed regions are shown magnified in b.i-iv. **c**-**d’**, Ex vivo culture procedure and representative images of *Gli1* (c) or Ki67 (d) staining and quantification (c’ and d’) in the indicated zones. n= 7 CTL, 5 EXP. Boxed regions are shown magnified on the sides. **e**, Proposed model of how CCN2 produced in specific cartilage zones limits *Gli1* expression in Pdgfra-derived chondrocytes during normal growth, whereas reduced CCN2 levels due to impaired growth in *Pan-Cart-p21^MOE^*limbs leads to unrestricted *Gli1* activation in this population, so that some of them become LLCPs.

To determine to what extent Pdgfra-lineage expansion was related to the expression of *Gli1* in cartilage cells, we analysed the overlap of tdTom (marking the Pdgfra-lineage) and *Gli1* expression in these cells at E17.5 (Fig. 6b). While the proportion of *Gli1*^+^ cells within the Pdgfra-lineage was increased from ∼40% in controls to nearly 60% in *Pan-Cart-p21^MOE^* samples (Fig. 6b’ squares), the proportion of Pdgfra-lineage cells within the Gli1^+^ population was slightly decreased from ∼70% to 55% (Fig. 6b’ circles). This suggests that the expansion of Gli1^+^ cells in the cartilage was due to cells residing both inside and outside the cartilage at the time of the insult. In summary, the number of Gli1^+^ chondrocytes– belonging to both the Pdgfra-lineage and the non-Pdgfra-lineage–was increased, unlike the Gli1^−^ Pdgfra- lineage, which remained the same in *Pan-Cart-p21^MOE^*as compared to controls (Fig. 6c).

Although *Pdgfra* mRNA levels were somewhat decreased in *Left-Cart-p21^MOE^* cartilage as compared to controls (Ext. Data Fig. 5c), we reasoned that chondrocytes that had recently formed from Pdgfra^+^ cells in response to cell-cycle arrest would retain PDGFRA protein for some time. Therefore, to further explore Gli1^+^ chondrocytes as derivatives of the Pdgfra-lineage, we analysed PDGFRA protein expression within and outside the Gli1-lineage at E14.5, i.e., shortly after p21 induction (Fig. 6d). Indeed, the number of PDGFRA^+^ cells was increased within the Gli1-lineage cartilage in *Pan-Cart- p21^MOE^* as compared to controls (Fig. 6d’ circles). In fact, the population that expanded the most in the *Pan-Cart-p21^MOE^* vs. control cartilage was the PDGFRA^+^ Gli1-lineage one (Fig. 6e), suggesting that this population is especially involved in the repair. In summary, our data suggests that Pdgfra^+^ cells (mostly located outside the cartilage) become Gli1^+^ CPs and progressively increase their contribution to cartilage in *Pan-Cart-p21^MOE^*limbs as compared to controls, contributing to the compensation. Lastly, we examined the behaviour of the Pdgfra-lineage in long-term normal growth. Tracing of fetal Pdgfra^+^ cells with the RGBow reporter during normal growth led to very few long columns being labelled at P30 (Fig. 6f, compare with 3c’), with most of the few long columns being restricted to the periphery of the growth plate (Ext. Data Fig. 7c-d’). This suggests that Pdgfra^+^ cells do not give raise to a significant number of Gli1^+^ LLCPs unless the cartilage is challenged. This begged the question of what the molecular changes triggered by the challenge were.

### CCN2 impairs Gli1 expression and proliferation in chondrocytes, and is downregulated in Cart-p21 samples

To determine how the *Pan-Cart-p21^MOE^* challenge triggers *Gli1* upregulation, we performed another transcriptomic approach, but this time shortly after challenge induction, separating Gli1-lineage and non- Gli1-lineage cells (Ext. Data Fig. 8a). Pregnant females carrying *Gli1^CreER/+^; R26^LSL-tdTom/+^; Tigre^TRE-p^*^21/+^ embryos (with or without *Col2a1-rtTA*) were provided with Dox at E12.5 and TM at E13.5, and embryos were collected at E14.5. The whole hindlimb was processed for single-cell isolation and 10x Chromium library preparation, multiplexed by sex (n=4 Exp and 4 Ctl samples, limbs from 2-3 embryos were pooled per sample, see Online Methods). Coarse clustering was initially generated and further curated refined manually (Ext. Data Fig. 8b, b’). Interestingly, in the coarse clustering, the two resting chondrocyte subpopulations formed a continuum with the cells showing markers of the perichondrial groove of Ranvier, a niche that hosts cells that can make contribution to cartilage^6^, supporting the connection between perichondrium and cartilage lineages. To identify gene signatures changed in *Cart-p21* limbs, we performed pseudobulk differential gene expression analysis–which minimises false discovery rates^24^–via edgeR (Ext. Data Fig. 8c). Besides the expected cell-cycle related genes (e.g. histone- encoding), we noticed changes in genes related with chondrocyte activity and response to stress, such as *cellular communication network factor 2* (*Ccn2*), also known as *connective tissue growth factor*^25–27^, which was downregulated in *Pan-Cart-p21^MOE^* limbs (see Violin plot and Gene Ontology analysis in Ext. Data Fig. 8c, d). We complemented these results by analysing cell-cell communication via MultiNicheNet^28^ in the E16.5 snRNA-seq and E14.5 scRNA-seq datasets. In all cases, *Ccn2*-mediated communication was significantly downregulated in experimental samples (Ext. Data Fig. 8e-f’). We confirmed this using *Ccn2* hybridisation chain reaction (HCR) on *Left-Cart-p21^MOE^*, *Pan-Cart-p21^MOE^* and control samples (Fig. 7b, b’, Ext. Data Fig. 8g, g’). These results were interesting considering our previous study in which we showed that overexpression of *Ccn2* in the left cartilage (*Pitx2-Cre/+; Col2a1-tTA/+; Tigre^Dragon-Ccn^*^2^*^/+^* model, Ext. Data Fig. 9a) impairs bone growth^15^. Using archival samples of this model, we observed that the left hypertrophic zone was extremely reduced in size (Ext. Data Fig. 9a’), and that proliferation in the resting zone (Ki67^+^ cells) was significantly downregulated in the left cartilage as compared to the right one or control littermates (Ext. Data Fig. 9b, b’).

Given the results above, and that CCN2 has been proposed to orchestrate various signalling pathways to promote harmonized skeletal growth^29^, we hypothesised that CCN2 interferes with *Gli1* activation in CPs, so that downregulation of *Ccn2* is what allowed the expansion of Gli1^+^ cells in the *Pan-Cart- p21^MOE^* models. To test this possibility, we cultured fetal femurs of control and *Pan-Cart-p21^MOE^* fetuses ex vivo (see Online methods) and treated them with human CCN2 (right femur from each embryo) or Vehicle (left femur from each embryo) for ∼2d (Fig. 7c). As predicted, *Gli1* expression was found downregulated in treated samples–both Ctl and *Pan-Cart-p21^MOE^*–(Fig. 7c’). Moreover, Ki67 immunostaining was used to detect proliferative cells, which were found to be diminished in treated samples (Fig. 7d, d’). In summary, we concluded that the expansion of the Gli1-lineage, and the compensatory proliferation triggered by cartilage cell-cycle arrest, were in part due to decreased CCN2 levels.

## Discussion

Elucidating the origin and regulation of the LLCPs is crucial to understanding the evolution of limb size and proportions, the factors that influence human height and the potential causes and treatments of skeletal dysplasias. In mice, postnatal CPs have been described that can self-renew and give rise to long columns of chondrocytes for most or the whole growth period^20,30,31^. However, given that at fetal and perinatal stages only short columns are generated^7^, it was unclear whether LLCPs are already present (albeit inactive) in the early cartilage, or they are formed later. In either case, we hypothesised that the system likely evolved with built-in robustness, such that transient developmental challenges may reveal regulatory details of the process. Following this approach, we found that fetal Gli1-derived chondrocytes expand in response to cell-cycle arrest in the cartilage and are required to compensate for this challenge (Figs. 1, 2). We further found that Gli1^+^ cells give rise to LLCPs during normal growth (Fig. 3). However, while we showed that the early cartilage anlage already contains Gli1^+^ quiescent CPs, our results also suggested that Gli1^+^ cells outside the initial cartilage anlagen contribute to the LLCP pool later, at least until birth (Fig. 4). Of note, chondrocytes arising from the Gli1-lineage in the last gestational week were shown to be critical for acquiring normal bone length in the hindlimb (Fig. 5). However, while bone length was significantly affected when Gli1-derived chondrocytes were ablated, the effect was somewhat mild. This was probably due to compensation after Dox clearance. While we tried to test this possibility by providing Dox postnatally, the rtTA that we used in our models did not get sufficiently activated.

Interestingly, our single-cell RNA-seq analyses identified a continuous cluster including two resting chondrocyte populations and one of cells expressing markers of the groove of Ranvier (Ext. Data Fig. 8b), with one of the resting populations within the Gli1^high^ subset being expanded at the expense of the groove-of-Ranvier one (Ext. Data Fig. 10). It is important to note, however, that this approach is limited because the Gli1 lineage had to be separated by FACS instead of bioinformatically identified, because in the 10x approach, which is 3’ biased, both the tdTom and the p21 transgenes yield reads mapping mostly to WPRE, the RNA-stabilising sequence used in both alleles (Fig. 1b).

As per the origin of the Gli1^+^ CPs, our results suggested that some chondrocytes derive from stromal Pdgfra^+^ cells (Fig. 6) revealing a previously unknown source of chondrocytes. Moreover, we showed that these Pdgfra^+^ cells do not give rise to Gli1^+^ LLCPs during normal growth, but that they can become *Gli1^+^* (and maintain PDGFRA expression at least briefly) in response to cartilage challenge. It remains to be determined whether there is a chemoattractant signal emanating from the cartilage, capable of recruiting Pdgfra^+^ cells. Such a signal could have therapeutic potential if it could be exogenously provided to stimulate CP formation in situ, in a minimally invasive manner. In summary, our data supports a model in which external cells always contribute to the cartilage, but mostly forming short-lived CPs, unless they switch their behaviour in response to a growth challenge.

Lastly, regarding the signalling changes operating in the challenged limbs, differential gene expression and cell-cell communication analyses showed that CCN2 levels were decreased in response to cartilage cell-cycle arrest (Fig. 7). Intriguingly, *Ccn2* downregulation also happened in the right limbs of the *Left- Cart-p21^MOE^* model (Fig. 7b, Ext. Data Fig. 8f). Since we have previously shown that both left and right limbs are equally affected in length in the *Left-Cart-p21^MOE^* model^11^, we propose that CCN2 levels correlate with the overall growth output during fetal bone growth. Moreover, here we showed that excess CCN2 impairs *Gli1* expression, chondrocyte proliferation and growth plate cytoarchitecture (Fig. 7 and Ext. Data Fig. 9), leading us to propose a model in which CCN2 acts as a negative feedback signal (Fig. 7e). In this model, CCN2 levels reflect the growth status of the cartilage, such that reduced growth output leads to decreased CCN2 and thus increased number of Gli1^+^ CPs, therefore unleashing long-term bone growth potential. One limitation of this model is that *Gli1* upregulation was mostly detected in the ExpL limb (Fig. 1a’), suggesting that CCN2 downregulation is necessary but not sufficient to derepress *Gli1* expression. Future studies will focus on elucidating additional factors that could be used to convert stromal cells in Gli1^+^ LLCPs, which could provide new avenues to stimulate cartilage regeneration in situ.

## Methods

### Animal models

The use of animals in this study was approved by the Monash Animal Research Platform animal ethics committee at Monash University.

### Unilateral p21 misexpression model

The *Pitx2-ASE-Cre* (aka *Pitx2-Cre*) mouse line, obtained from Prof. Hamada^9^, was crossed with the *Col2a1-rtTA* mouse line^10^, and then inter-crossed to generate *Pitx2-Cre/Cre; Col2a1-rtTA* mice. In some experiments, a *Col2a1-eCFP* allele^32^ was also included (a kind gift from David Rowe). Genotyping was performed as described previously^11^. The *Tigre^Dragon-p^*^21^ mouse line was previously described^11,15^. Experimental and control animals were generated by crossing *Pitx2-Cre/Cre; Col2a1-rtTA/+* females with *Tigre^Dragon-p^*^21^*^/Dragon-p^*^21^ males (i.e. homozygous for the conditional misexpression allele). The separation of control and experimental animals was based on the rtTA genotype. Pregnant females were administered doxycycline hyclate (Sigma) in water (1 mg/kg with 0.5% sucrose for palatability) from E12.5 until collection (or until birth). The day of vaginal plug detection was designated as E0.5, and E19.5 was referred to as P0.

### Multi-colour lineage tracing

*Gli1^CreER/CreER^* female mice were time-mated with *R26R^RGBow/RGBow^* male mice. The day of vaginal plug detection was referred to as E0.5, and E19.5 as P0. Pregnant females were administered Tamoxifen (Sigma, stock at 20 mg/ml in corn oil) by oral gavage (180 µg/g) at E12.5 (Experiment1) or E13.5 (Experiment2). Mouse embryos were collected at E14.5, E16.5, P0, P1, P14, P30 (Experiment1) or E14.5,E15.5, E17.5,P0,P1,P14,P30 (Experiment2). Limbs were dissected and fixed in 4% paraformaldehyde for 2d at 4°C. Decalcification was performed on P14-P30 samples by immersing them in 0.45M ethylenediaminetetraacetic acid (EDTA) in PBS from 10-14 days at 4°C.

### Pulse of nuclear GFP expression in cartilage

*Gli1^CreER/CreER^; Tigre^TRE-nGL/TRE-nGL^* female mice were time-mated with *Col2a1-rtTA/+*; *R26^LSL-tdTom/LSL-tdTom^* male mice. Experimental set 1: Pregnant females were administered Dox in drinking water and Dox food (Specialty feeds SF08-026, 600mg/Kg) from E12.5-E13.5. Tamoxifen was administered by oral gavage (180µg/g) at E13.5. Mouse embryos were collected at E14.5,E17.5,P0,P14,P21. Experimental set 2: like set 1, except that Dox was given from E17.5-E18.5, and offspring was collected at P0, P14,P21 and P30. Experimental set 3: Like set 2, except that Dox was given from E12.5-E18.5, and offspring collected at P14. Limbs processed as above.

### Lineage tracing combined with p21 expression in all cartilage elements

*Gli1^CreER/CreER^* females^33^ were crossed with *Col2a1-rtTA/+; Tigre^TRE-p^*^21^*^/TRE-p^*^21^*; R26^LSL-tdT/LSL-tdT^* males (described in ^15,17^). Pregnant females were given Dox food and water, plus tamoxifen (180 µg/g orally), at the stages described in the main text. To trace other lineages, *Pdgfra^CreER^* females^23^ (a kind gift from Brigid Hogan) were used instead of *Gli1^CreER^* ones (tamoxifen 100 µg/g). *Pdgfra^H2B-EGFP^* mice^34^ were a gift from Philippe Soriano.

### Unilateral *Ccn2* misexpression model

The *Pitx2-Cre* mouse line was crossed with the *Col2a1-tTA* one that we previously described^11^, and then inter-crossed to generate *Pitx2-Cre/Cre; Col2a1-tTA* mice. The *Tigre^Dragon-Ccn^*^2^ mouse line was previously described^15^. Experimental and control animals were generated by crossing *Pitx2-Cre/Cre; Col2a1-tTA/+* females with *Tigre^Dragon-Ccn2/Dragon-Ccn2^* males (i.e. homozygous for the conditional misexpression allele). The separation of control and experimental animals was based on the tTA genotype. Pregnant females were not given Doxycycline, to allow for tTA activity from the onset of chondrogenesis.

### Generation of G4 mouse embryonic stem cells with Tigre docking site

a TIGRE docking site targeting vector (a gift from Hongkui Zeng via Addgene plasmid #61580; http://n2t.net/addgene:61580; RRID:Addgene_61580) was electroporated by Monash Genome Modification Platform into G4 mESCs (a gift from Andras Nagy) to generate an ESC line (MG36) with a docking site in the *Tigre* (*Igs7*) locus, amenable for Flp-mediated cassette exchange. 192 individual clones were picked after 6 days of G418 selection. Out of the 96 analysed clones, 21 were positive by Southern hybridisation with both 5’ and 3’ probes. After reanimation, these clones were analysed again by Southern hybridisation, 8 positive clones were further characterised by Southern hybridisation using internal neo probe (3 restriction digestions); 7 clones (#9, 13, 17, 32, 56, 61, 62) showed single integration. After sequencing, all positive clones were found to have the correct sequence at F5 and F3 mutant FRT sites. 4 clones were karyotyped: clones #56 and 62 are 100% euploid metaphase, whereas clone#32 was 87% euploid, and clone#9 80% euploid.

### Generation of Tigre^TRE-nucGreenLantern^

AR157 vector, containing a TRE-H2B-GreenLantern and a PGK-Hygro resistance split gene (N-half) flanked by FRT sites was generated by VectorBuilder (Exp Data Fig. 6a). The vector also contained a Polylox cassette^35^ downstream of the GreenLantern, but this feature was not used in this study. This vector was then used by Monash Genome Modification Platform for Flp-mediated cassette exchange (Exp Data Fig. 6a) into clones #56, 62 and 32 of *Igs7-* targeted G4 ES cells (MG36, see above). 30µg pCAG-FlpO-IRES-Puro + 10µg AR157, mixed with 0.8ml of ESC suspension at ∼1x10^7^/ml and transferred to a 4mm cuvette. After incubation at RT for 5 min, electroporated by applying a single pulse (an exponential decay pulse at 250V, 500µF capacitance or a square wave pulse at 250V, 5ms duration) followed by 5min at RT, then added into a 10-cm dish pre-coated with MEFs. Two electroporation (one exponential decay pulse and one square wave pulse) per clone was performed. 1 day after transfection, transfection efficiency was checked under the fluorescent microscope, and drug selection was started (150µg/ml hygromycin). At 4 days after transfection, selection was changed to hygromycin 100µg/ml. At 6 days after transfection, we picked up 4 clones from each parental clone (#56-1 to 4, #62-1 to 4, #32-1 to 4) to MEF-coated 24-well plates. These clones were genotyped by 5’- GT PCR, 3’- GT PCR and TRE-GL specific PCR (see below). All clones were correctly targeted (Ext. Data Fig. 6b).

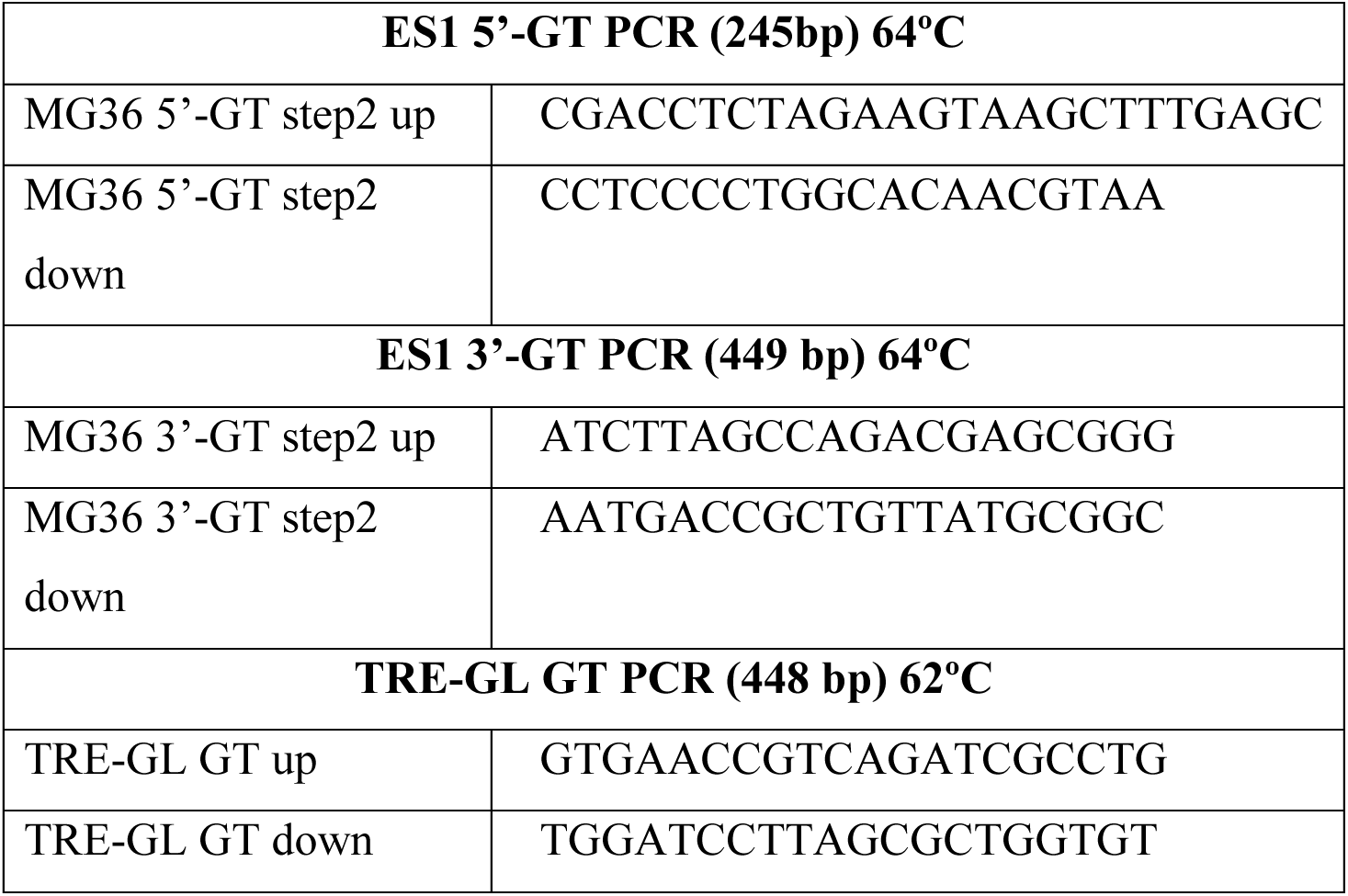

Clones #62-1, 62-2, 56-1 and 56-2 were used for two sessions each of blastocyst microinjection. Males ET6 (clone 62-1, 85% chimera), ET28 (clone 62-2, 85% chimera), ET51 (clone #56-1, 80% chimera), ET59 (clone #56-2, 80% chimera) were selected for breeding to next generation. One of these lines was crossed with *Gli1^CreER^*animals to establish the *Gli1^CreER^; Tigre^TRE-nGL^* lines. Six additional highly chimeric males were kept as back-up breeders.

### X-gal staining

For enzymatic detection of β-galactosidase activity, the frozen sections were postfixed 5 min with 4% paraformaldehyde (PFA, Electron Microscopy Sciences) in PBS at RT. After PBS washes, the sections were incubated 2 x 5 min with X-gal buffer (2 mM MgCl_2_, 0.02% NP40 and 0.05% deoxycholate in PBS 0.1 M pH 7.4) and then overnight at 37°C in X-gal reaction buffer (20 mg/ml X-gal, 5 mM K_4_Fe(CN)_6_ and 5 mM K_3_Fe(CN)_6_ in X-gal wash buffer). After PBS rinses, the sections were postfixed 10 min in 4% PFA and PBS-rinsed again. The sections were then counterstained with Nuclear Fast Red 0.005% for 15 min, serially dehydrated, incubated 3 x 1 min with xylene, and cover-slipped using DPX mountant (Fisher).

### Single-nuclei and single-cell RNA-seq and analysis

#### Single-nuclei RNA sequencing of unilateral p21 model

*Pitx2-Cre/Cre; Col2a1-rtTA/+; Col2a1-eCFP/+* females were crossed with *Tigre^Dragon-p^*^21^*^/Dragon-p^*^21^ males and given Dox (1mg/ml in drinking water with 0.5% sucrose) at E12.5. The knee region (obtained via orthogonal cuts from the hypertrophic region of the distal femur to the hypertrophic region of the proximal tibia in skinned hindlimbs) of E16.5 Left and Right limbs from uCtl and *Left- Cart-p21^MOE^* samples (2 embryos of each genotype) were collected in DMEM with 10% DMSO and 5% calf serum, stored in a MrFrosty (ThermoFisher) at 4°C for 10 min and then transferred to -80°C overnight. Tissue lysis and nuclei isolation were done as described by L.G.M. in https://www.protocols.io/view/frankenstein-protocol-for-nuclei-isolation-from-f-5jyl8nx98l2w/v2, in two batches of four samples each (left and right knees from one control and one experimental embryo). Once ∼5,000 nuclei per sample were collected, we proceeded immediately with the 10x Genomics Single Cell Protocol (https://www.10xgenomics.com/support/single-cell-gene-expression/documentation/steps/library-prep/chromium-single-cell-3-reagent-kits-user-guide-v-3-1-chemistry, minimizing the time between nuclei preparation/sorting and chip loading. One chip per batch was used (4 channels in each case, to load left and right samples from one control and one experimental embryo). The libraries were sequenced twice on Novaseq (800 million clusters/read pairs each time, which when combined amounted to ∼40,000 reads per nuclei across the 8 samples).

Following sequencing, fastq reads were mapped against a custom reference that combined Mus musculus (GRCm38) with transgenic genes (tdTomato, ECFP, and WPRE) using CellRanger (v7.1.0). The resulting aligned reads were analysed via R (4.0.2). Seurat (v.4.2.1) and R/Bioconductor SingleCellExperiment packages were used to generate count matrices. For each sample, the mean of unique molecular identifiers (UMIs) (nCounts) was calculated and used as a threshold to exclude empty droplets. Filtering was manually done according to the nCount and number of genes. DoubletFinder (v2.0.3) was used to remove the expected doublets in each sample. Harmony R package (v0.1.0) was also used to mitigate remove batch effects, and subsequently the data were consolidated into a single Seurat object.

To identify the appropriate resolution, Clustree v 0.5.0 and Cluster Correlation analysis were used. Visual representation of cluster assignments for individual cells was achieved within a reduced-dimensional space using UMAP. To pinpoint the differentiation marker genes in the specific cluster as compared to the remaining clusters, Wilcoxon Rank-Sum Test was used. This analysis was conducted using the “FindAllMarkers” function within Seurat (v.4.2.1), p-value of less than 0.05 and only.pos = T, and a log-fold change threshold of 0.1. To identify the cell type annotation, the clusters were mapped with published mouse single cell hindlimb data^36^. In addition, previously reported cellular biomarkers and Gene Ontology (GO) annotations were utilized to annotate the cell clusters. The top 100 marker genes were utilized to enrich Gene Ontology (GO) terms through the ClusterProfiler package (version 3.14.3). To identify and confirm the cell type, GSEA comparison was also performed using SCP package (V. 0.5.1).

### Single Cell RNA-sequencing

#### Sample preparation, collection, and sequencing

Female *Gli1*^CreER/CreER^ mouse was crossed with *Col2a1-rtTA; Tigre^TRE-p^*^21^*^/TRE-p^*^21^; *R26^LSL-tdT/LSL-tdT^* male. At E12.5, the pregnant mouse was administered by doxycycline in water (1 mg/ml with 0.5% sucrose) and food, and Tamoxifen was orally gavaged (120 μg/g) at E13.5. At E14.5, the embryos were carefully dissected. Initially, the sex and rtTA genotyping were determined using PCR-based quick genotyping and hindlimbs were then meticulously dissected.

Afterwards, the tissue underwent treatment with Accumax in a rocker at 31°C for 25 minutes (Ext. Data Fig. 3a) and every 10 min, the samples were gently shaken. The supernatant was carefully transferred into a tube and placed on ice for preservation, while the cartilaginous portions were exposed to Collagenase B (5 mg/ml; Sigma) for ∼90 min at 37°C in a water bath (pipetted every 20 min). To stop collagenase activity, we then added 700 µl of cold DMEM+20% FBS for each tube. We then incorporated outer and inner supernatants, and the samples were resuspended in 20 volumes of 1x RBC lysis buffer (including NH_4_Cl, NaHCO_3_, and EDTA) and incubated for 10 min RT on a rocker. The cells were then centrifuged at 500 xg for 5 min at 4°C and the cell pellet was then suspended in 900 µl of PBS + 0.05% BSA. After re-washing, and the cells were resuspended in 400 ul of PBS + 0.05% BSA. At the end, the cell suspension was filtered through 40-μm Flowmi® Cell Strainers and cells were stained with DAPI (0.05 µg/ml) and transferred into Falcon™ Round-Bottom Polypropylene (Fisher Scientific). The tdT+ and tdT- cells were sorted using flow cytometry (nozzle size of 100 µm, operating at a PSI of 20 and a BOP of 223).

In total, two experimental males, two control males, two control females, and only one experimental female were utilized. The cells were counted by an automated cell counter (ThermoFisher) using trypan blue staining and all passed at least 90% cell viability. Then, the cell suspensions were centrifuged at 500 xg for 5 min at 4°C to decrease the final volume. The samples (S) were then multiplexed based on <u>sex and conditions</u>. Specifically, for S1, EM^tdt+^ (Experimental Male) and CF^tdt+^ (Control Female) cells were multiplexed, totalling 7,500 cells. For S2, EM^tdt-^ and CF^tdt-^ cells were multiplexed, amounting to 25,000 cells (at a concentration of 1000 cells/µl with a volume of 25 µl). In S3, EF^tdt+^ and CM^tdt+^ cells were multiplexed, totalling 5,700 cells. Finally, in S4, EF^tdt-^ and CM^tdt-^ cells were multiplexed, comprising 24,000 cells. After that, the process was carried out according to the 10X Genomics protocol (https://www.10xgenomics.com/support/single-cell-gene-expression/documentation/steps/sample-prep/single-cell-protocols-cell-preparation-guide).

The libraries were prepared using Chromium Next GEM Single Cell 3’ Reagent Kits v3.1 (10X Genomics) according to manufacturer’s directions and then sequenced on Illumina (∼833 million clusters/read pairs each time) with a length of 100 bp. All samples were passed the quality of cDNA (refer to the GSE File).

### scRNA-seq preprocessing

The fastq files were aligned against a custom reference comprising *Mus musculus* (GRCm38; obtained from 10X Genomics Ref.) along with transgenic genes (tdTom, CreER^T2^, and WPRE) using CellRanger (v7.2.0). In total, more than 98.2% of Reads Mapped to Genome and 25,190 genes were detected as well as transgenes. To eliminate contaminated cells with ambient RNA, CellBender V 0.3.0 was used. The expected number of genuine cells and total droplets were estimated based on the CellRanger output, using the Raw_feature_bc_matrix.h5 file as input. Identified unique molecular identifiers (UMIs) were then mapped and subset from the Seurat object generated by the filtered feature barcode matrix file. DoubletFinder (v2.0.3)^37^ was then used to remove expected doublets in each sample, with pK calculated separately for each sample. Then the cells with low nCount_RNA and nFeature_RNA were filtered out.

### Testing for differences in cell type proportions

To analyse the distinct cellular compositions among conditions, the propeller test available in the speckle R (v.0.0.1, accessible at https://github.com/Oshlack/speckle) was employed. The groups were classified with a false discovery rate (FDR) of ≤0.1 as indicative of notable variations in cell types.

### MultiNicheNet

To investigate cell-to-cell communication, MultipleNichenetR (version 1.0.0)^38^ was used. The Seurat object was first converted to a SingleCellExperiment (SCE), and all genes were mapped using the alias_to_symbol_SCE function to "mouse." A cutoff point of min_cells = 10 was used, and differentially expressed genes were calculated using the get_DE_info function. The following parameters were applied: logFC_threshold = 0.5, p_val_threshold = 0.05, fraction_cutoff = 0.05, and empirical_pval = TRUE. To achieve more robust results and minimize cluster dropouts caused by low number of cells, some subclusters were merged. Ligand-receptor interactions were identified via the “multi_nichenet_analysis” function, and the results were visualized using MultiNicheNetR’s built-in functions^38^.

#### SCENIC analysis

SCENIC (version 1.2.4)^39^ was used to assess gene regulatory network activities in each cell population. For snRNA-seq data, pySCENIC^40^ was employed and then data manually converted to SCENIC in R. A co-expression network was generated using the GENIE3 function, and the Area Under the Curve (AUC) was calculated with AUCell. The AUC values were normalized to a range of 0 to 1 for each regulon and regulon scores were obtained using the ’get_AUC’ function. To identify significant changes in regulon activity across conditions, t-tests and Wilcoxon tests were conducted for each cluster in the relevant condition.

### EdU incorporation and detection

A solution of 5-Ethynyl-2’-deoxyuridine (EdU) was prepared at 6 mg/ml in phosphate-buffered saline (PBS). This solution was administered at a dose of 60 µg/g, subcutaneously (s.c) for pups and 30 µg/g intraperitoneally (i.p.) for pregnant females, 1.5 hours before euthanizing the mice (or 2 days before, for pulse-chase experiments). To detect EdU, a click chemistry reaction with fluorescein-conjugated azide (Lumiprobe #A4130) was performed once the immunohistochemistry and/or TUNEL staining were completed on the same slides. Briefly, the working solution was prepared in PBS, adding CuSO_4_ (Sigma # C1297) to 4 mM, the azide to 0.4 µM and freshly-dissolved ascorbic acid (Sigma #A0278) to 20 mg/ml, and incubating the sections for 15 min at room temperature in the dark, followed by PBS washes.

### Sample collection and histological processing

Mouse embryos were euthanized using hypothermia in cold PBS, while mouse pups were euthanized by decapitation. Upon collection of the embryos or pups, the limbs (including full tibiae and/or femora) were carefully dissected out in cold PBS, skinned, and fixed in 4% paraformaldehyde (PFA) for 2 days at 4°C. Samples of P1 or younger were not subjected to decalcification. For P3, P5, and P7 samples, decalcification was performed by immersing the specimens in 0.45M ethylenediaminetetraacetic acid (EDTA) in PBS for 3, 5, and 7 days, respectively, at 4°C. Following several washes with PBS, the limb tissues were cryoprotected in PBS containing 15% sucrose and then equilibrated in 30% sucrose at 4°C until they sank. The hindlimbs were then oriented sagittally, facing each other, with the tibiae positioned at the bottom of the block (closest to the blade during sectioning) and embedded in Optimal Cutting Temperature (OCT) compound using cryomolds (Tissue-Tek). The specimens were frozen by immersing them in dry-ice-cold iso-pentane (Sigma). Serial sections with a thickness of 7 µm were collected using a Leica Cryostat on Superfrost slides. The sections were allowed to dry for at least 30 minutes and stored at -80°C until further use. Prior to conducting histological techniques, the frozen slides were brought to room temperature in PBS, and the OCT compound was washed away with additional rounds of PBS rinses.

### Ex vivo culture

The culture medium was prepared by mixing 200 μL penicillin/streptomycin (10,000 U/mL) with 48.9 mL high-glucose DMEM, 500 μL Glucomax, 100 mg BSA, 2.5 mg ascorbic acid, and 400 μL β-glycero-phosphate, then filtered through a 0.2 μm filter. E14.5 embryos were dissected and transferred to pre-warmed media under a tissue culture hood. Each hindlimb was placed in cold Accumax and then digested at 31°C for at least 35 minutes. The samples were then put on ice, and the tibia and femur were dissected individually. Limbs were cultured in 48-well plates with 500 μL of media at 37°C in a 5% CO_2_ incubator for approximately 2 hours before initiating treatment. For the treatment, human recombinant CCN2 (Peprotech, 17840573) was used. One limb from each pair was treated with 100 ng/mL of CCN2 + Doxycycline, while another set received Doxycycline + Vehicle (1% BSA in Saline). After 2 days, the limbs were treated with 10 µM EdU for 1h before fixation in PFA.

### Micro-CT and bone length analysis

The micro-CT and bone length analysis was as previously described^41^. Briefly, samples were retained and fixed in 4% PFA as residual tissues from other experiments in the Rosello-Diez lab. Whole femora and humeri were scanned using a Siemens Inveon PET-SPECT-CT small animal scanner in CT modality (Monash Biomedical Imaging). The scanning parameters included a resolution of 20 and 40 µm, 360 projections at 80 kV, 500 µA, 600 ms exposure with a 500 ms settling time between projections. Binning was applied to adjust the resolution with 2 x 2 for 20 µm scans and 4 x 4 for 40 µm scans. The acquired data were reconstructed using a Feldkamp algorithm and converted to DICOM files using Siemens software. For the analysis and bone length measurements, Mimics Research software (v21.0; Materialize, Leuven, Belgium) equipped with the scripting module was utilized to develop the analysis pipeline^41^.

### Immunohistochemistry and TUNEL staining

For antigen retrieval, the sections were subjected to citrate buffer (10 mM citric acid, 0.05% Tween 20 [pH 6.0]) at 90°C for 15 minutes. Afterward, the sections were cooled down in an ice water bath, washed with PBSTx (PBS containing 0.1% Triton X-100). To perform TUNEL staining, the endogenous biotin was blocked using the Avidin/Biotin blocking kit (Vector #SP-2001) after antigen retrieval. Subsequently, TdT enzyme and Biotin-16-dUTP (Sigma #3333566001 and #11093070910) were used according to the manufacturer’s instructions. Biotin-tagged DNA nicks were visualized using Alexa488- or Alexa647-conjugated streptavidin (Molecular Probes, diluted 1/1000) during the incubation with the secondary antibody.

For immunohistochemistry staining, sections were incubated with the primary antibodies prepared in PBS for either 1.5 hours at room temperature or overnight at 4°C (see list of antibodies below). Following PBSTx washes, the sections were incubated with Alexa488-, Alexa555-, and/or Alexa647- conjugated secondary antibodies (Molecular Probes; diluted 1/500 in PBSTx with DAPI) for 1 hour at room temperature. After additional PBSTx washes, the slides were mounted using Fluoromount™ Aqueous Mounting Medium (Sigma). The antibodies used, along with their host species, vendors, catalogue numbers, and dilutions, were as follows: mCherry (goat polyclonal, Origene Technologies #AB0040-200, diluted 1/500), p-S6 (rabbit polyclonal, Cell Signaling Technology #2211S, diluted 1/300).

### In situ hybridization

To perform *in situ* hybridisation, sections were fixed in 4% PFA for 20 minutes at room temperature, washed in PBS, and treated with 4µg/ml Proteinase K for 15 minutes at 37°C. After washing in PBS, the sections were refixed with 4% PFA, followed by treatment with 0.1N pH8 triethanolamine (Sigma #90279), 0.25% acetic anhydride (Sigma #320102) for 10 minutes at room temperature. Subsequently, the sections were washed in PBS and water and incubated with prehybridization buffer (50% formamide, 5x SSC pH 5.5, 0.1% 3-[(3-Cholamidopropyl)dimethylammonio]-1-propanesulfonate (CHAPS), 0.05 mg/ml yeast tRNA, 0.1% Tween 20, 1x Denhardt’s) at 60°C for 30 minutes. The sections were then incubated with 1µg/ml preheated riboprobes and subjected to hybridization at 60°C for 2 hours. Post-hybridization washes were performed using post-hybridisation buffer I (50% formamide, 5x pH 5.5 SSC, 1% SDS) and II (50% formamide, 2x pH 5.5 SSC, 0.2% SDS) preheated at 60°C for 30 minutes, respectively. The sections were then washed with maleic acid buffer (MABT: 100mM maleic acid, 150mM NaCl, 70mM NaOH, 0.1% Tween 20) and blocked with 10% goat serum, 1% blocking reagent (Roche #11096176001) in MABT at room temperature for 30 minutes. Next, the sections were incubated overnight at 4°C with anti-digoxigenin-AP (Sigma #11093274910) diluted 1/4000 in MABT with 2% goat serum and 1% blocking reagent. After several MABT washes, the sections underwent incubation with AP buffer (0.1M Tris-HCL pH 9.5, 0.1M NaCl, 0.05M MgCl_2_, 0.1% Tween 20), with the second one containing 1mM levamisole and then developed colour using BM purple (Roche #11442074001) at 37°C. Following a wash in PBS, the sections were fixed for 10 minutes in 4% PFA, counterstained with Nuclear Fast Red (Sigma #N3020) at room temperature for 10 minutes, and rinsed in water. Dehydration was performed by passing the sections through 70%, 90% ethanol, absolute ethanol, and xylene. Finally, the sections were mounted with DPX (Sigma #100579).

### Hybridisation chain reaction (HCR)

Slides were thawed for 20 min in RNAse-free PBS (DPBS; GIBCO, Invitrogen, LOT: 866200), washed twice with DPBS, and fixed in ice-cold 4% paraformaldehyde for 15 min at 4°C. Samples were then dehydrated in 50%, 75%, and 100% ethanol for 5 min each at room temperature. After washing with DPBS, slides were placed in a pre-warmed humidified chamber, and 200 µL of probe hybridization buffer was added for 10 min. Probes (95 µL, 16 nM) were then applied and incubated for > 12 hours at 37°C.

To remove excess probes, slides were washed sequentially with 75%, 50%, and 25% probe wash buffer (24 mL formamide 100% with 20 mL 20x SSC, 1,440 µl of Citric acid 0.5 M, 80 µl Tween20, 200 µl 20 mg/mL heparin, and fill up to 80 mL with ultrapure water) in 5x SSCT, followed by 100% 5x SSCT, each for 15 min, then immersed in 100% 5x SSCT for 5 min at room temperature. Amplification was initiated by adding 200 µL of amplification buffer for 30 min at room temperature. Hairpins h1 and h2 (6 pmol each) were snap-cooled and mixed with amplification buffer. After removing the pre-amplification solution, 98 µL of the hairpin solution was added, and slides were incubated for at least 12 hours in a dark, humidified chamber. Slides were washed twice with 5x SSCT for 30 min and once for 5 min, then mounted with Fluoromount-G™ (Cat: 00495802, USA).

### Imaging

Sagittal sections of the limbs were captured, with a focus on the area between the distal femora and proximal tibiae. Typically, at least 2 sections per limb were analysed, although in most cases 4 sections were examined. In the case of cultured distal femora, frontal sections were used as they provided better identification of the different epiphyseal regions. To determine the boundaries of the resting zone (RZ), proliferative zone (PZ) and hypertrophic zone (HZ), morphological criteria were applied. The transition between round (resting) and flat (columnar) nuclei, forming an arch along the upper point of the grooves of Ranvier, was considered as the start of the PZ. On the other hand, the transition towards larger, more spaced nuclei (pre-hypertrophic) marked the end of the PZ. The point where the pericellular matrix exhibited a sharp reduction around enlarging chondrocytes was designated as the beginning of the HZ. The distal end of the last intact chondrocyte served as the endpoint of the HZ.

For imaging, bright-field and fluorescence images were acquired using a Zeiss inverted microscope (Imager.Z1 or Z2) equipped with Axiovision software (Zeiss/ZenBlue). Mosaic pictures were automatically generated by assembling individual tiles captured at 10x magnification for bright-field images or 20x magnification for fluorescence images.

### Image analysis and quantification

The regions of interest such as resting, proliferative and hypertrophic zones (RZ, PZ, HZ) and the secondary ossification centre (SOC) were identified from imaged sections of the multiple models (left and right; experimental and control proximal tibial cartilage). Consistent parameters such as brightness, contrast, filters and layers were kept the same for all the images in the same study.

### Cell count analysis

The RZ and PZ were identified and segmented from sections stained for DAPI, tdTomato, and TUNEL or EdU using macros on FIJI software. Details of the macros are available upon request. The number of cells in the region of interest (RZ & PZ) were measured using CellProfiler. Since in our hands CellProfiler did not work properly with more than 3 channels, a pipeline was generated to split the four channels into DAPI on one hand and the 3 other channels on the other hand. The two sets of images were run on CellProfiler separately. The CellProfiler pipelines are also available upon request.

### Statistical analysis

Statistical comparisons were performed using appropriate tests based on the experimental design. An unpaired t test was used for comparisons involving one variable and two conditions. Two-way ANOVA was utilized for comparisons involving two variables and two or more conditions. Non-parametric tests were selected when the assumption of normality could not be met. All statistical analyses were conducted using Prism10 software (GraphPad).

## Supporting information

Ext. Data Fig.

## Acknowledgments

We acknowledge the Monash Genomics and Bioinformatics Platform for their excellent technical help, as well as the Monash Genome Modification Platform, for their help with gene targeting and genotyping strategy (design and execution). We thank Timothy Semple for his help with library preparation for the snRNA-seq dataset, and Ricky Johnstone for providing access to bioinformatics expertise.

## Conflict of Interest

F.J.R. receives institutional support as a coinvestigator and subcontracted by the Peter MacCallum Cancer Centre for an investigator-initiated trial which receives funding support from Sanofi/Regeneron Pharmaceuticals. The rest of the authors declare that the research was conducted in the absence of any commercial or financial relationships that could be construed as a potential conflict of interest.

## Author Contributions

X.Q.: data acquisition and analysis, supervision; E.R.: data acquisition and analysis, figure preparation, manuscript editing; S.L.A.: data analysis, manuscript editing, supervision; K.K.V.: data acquisition and analysis, figure preparation; F.J.R.: data analysis; M.Z.: data analysis; L.G.M.: experimental design, data acquisition; C.H.H.: data analysis, manuscript editing; D.R.P.: tool generation, data analysis, supervision; A.R-D.: conceptualization, experimental design, data analysis, funding acquisition, supervision, figure preparation, manuscript drafting.

## Funding

HFSP CDA00021/2019-C (to A.R-D.) and NHMRC Ideas grant 2002084 (to A.R-D. and C.H.H.). The Australian Regenerative Medicine Institute is supported by grants from the State Government of Victoria and the Australian Government. The Novo Nordisk Foundation Center for Stem Cell Medicine, reNEW, is supported by a Novo Nordisk Foundation grant number NNF21CC0073729.

## Data Availability Statement

The single-nuclei and single-cell RNA-seq datasets have been deposited at the NCBI Sequence Read Archive (SRA) with accession numbers PRJNA1136579 and PRJNA1138445, respectively. These can be accessed via the Gene Expression Omnibus (GEO), with accession numbers GSE273540 and GSE273538, respectively.

## Extended Data Figure legends

**Extended Data Figure 1.** Possible origins of postnatal cartilage progenitors (CPs). **a**, Cartilage-resident short-lived CPs eventually become long-lived, due to intrinsic and/or extrinsic triggers. **b**, Precursors of long-lived CPs (Pre-LLCPs) already exist in the fetal limb as a separate population (in the cartilage and/or outside), and only become active/recruited postnatally, in response to cartilage maturation. RZ, PZ, HZ, resting, proliferative, hypertrophic zones; SOC, secondary ossification centre; PCh, perichondrium.

**Extended Data Figure 2. a**, Double-conditional model for inducible and reversible gene overexpression. TRE, Tetracycline-responsive element. WPRE, RNA stabilising sequence. **b**, Expected accelerated depletion of chondroprogenitors in response to mosaic p21 expression. **c**, Minor to no asymmetries are generated by P100 despite the overactivation of perinatal CPs.

**Extended Data Figure 3. a**, Schematic of the process for sample collection, processing of single-nuclei RNA sequencing and quality control in Seurat processing. **b**-**b’**, Dotplot graph illustrating the markers of each of the 32 initially identified clusters (b) and chondrocyte-related clusters only (b’). **c**, Heatmap of average gene expression of all clusters showing the accuracy of clustering. **d**-**d’**, UMAP of all identified clusters (d). The highly correlated clusters (lateral plate mesoderm-derived cells, LPM) are shown boxed by black rectangles on the heatmap (c), and subclustered in (d’). **e**-**e’**, Gli1+ cells in the LPM group (e) and distribution of LPM populations within Gli1^+^ cells in Ctl and Exp limbs (e’). * denotes a significant difference identified by Propeller.

**Extended Data Figure 4. a**, **a’**, % of p21^+^ nuclei found in the proximal tibia cartilage of Cart-p21 mice at the shown stages, with Dox at E12.5 (a) or E13.5 (a‘). p-values of multiple-comparisons tests after ANOVA are shown. **b**, The Gli1 lineage was followed in Control and Cart-p21 samples, and quantified inside and outside the cartilage (defined in Fig. 4c). **c**-**c’’**, Distribution of tdT^+^ cells in resting (RZ) and proliferative zone (PZ) of Ctl and *Pan-Cart-p21^MOE^* samples at E14.5 (c), E17.5 (c‘) and P0 (c’‘). **d**, **d’**, Idem, for P0 samples with TM-E12.5 and Dox at E13.5 (d) or E12.5 (d‘). In b-d’, p-values for multiple comparisons tests after 2-way ANOVA are shown.

**Extended Data Figure 5.** Expression of Gli1 (a, a’) and Pdgfra (b, b’) in the embryonic day (E)14.5 and postnatal day (P)1 hindlimb, as indicated. Fe, femur, Ti, tibia. Dashed lines delineate the cartilage. Some images were taken from GenePaint (a, b).

**Extended Data Figure 6. a**, Targeting strategy to generate ES cell lines carrying a Tet-responsive element (TRE)-controlled nuclear green lantern (H2B-GL) gene in the Tigre locus. Successful FlpO- mediated cassette exchange reconstitutes the Hygro-resistance gene. SA, splice acceptor. pA, poly A. Ins, insulator. FRT, flippase recognition target. WPRE, RNA-stabilising sequence from Woodchuck virus. PGK, constitutive mammalian promoter. **b**, Confirmation of initially picked clones (left) and subclones (right) by 5’/3’/TRE-GL specific PCR reactions. **c**, **c’**, Representative images (c) and quantification in the yellow region of interest (c’) of the single- and double-labelled cells, upon TM and Dox induction, as indicated. **d**, Quantification of Ki67^+^ (i.e., cycling) cells, both GL^+^ and GL^−^, in the RZ. Stages and Dox treatment are indicated. p-values for multiple comparisons test after 2-way ANOVA are shown.

**Extended Data Figure 7. a**-**b’**, Quantification of EdU in p21-negative chondrocytes of resting and proliferative zones (a, b, as indicated) and in cells outside the cartilage (a’, b’), at E14.5 (a-a’) and E17.5 (b-b’), distinguishing between Pdgfra-lineage and non-lineage cells. In (a-b’), p-values for multiple comparisons test after 2-way ANOVA are shown. **c**-**d’**, Tri-colour lineage tracing of Pdgfra^+^ cells, upon TM injection at E12.5 (c, d, boxed region amplified in c’) and E13.5 (d’). For the quantifications in d and d’, the average number of columns of each colour per sections was added up to obtain a total number of columns per section per sample.

**Extended Data Figure 8. a**, Design of the lineage-specific single-cell RNA-seq approach, multiplexed by sex. **b**, **b’**, Coarse (b) and fine (b’) clustering outcome of the curated sequencing data (see Online Methods). **c**, Violin plot showing expression of *Ccn2* in the different populations identified in the scRNA-seq data, for the 2 genotypes. CTL= CTN+CTP, EXP= ETN+ETP. **d**, Gene Ontology terms related to Ccn2, differentially enriched in EXP vs. CTL samples. **e**-**f**’, MultiNicheNet analysis of cell- cell communication in Proliferating Chondrocytes of the snRNA-seq (e) and the scRNA-seq dataset (f), as well as Resting chondrocytes of the scRNA-seq dataset (f’). In e, f-f’, Ccn2 is boxed. **g**, **g’**, Representative images (g) and quantification (g’) of *Ccn2* expression as determined by HCR. PZ, RZ, GoR: proliferative zone, resting zone, groove of Ranvier (separated from cartilage by dashed line). n= 5 Ctl, 5 *Pan-Cart-p21^MOE^*.

**Extended Data Figure 9. a**, **a’**, Mouse model of unilateral, cartilage-targeted, *Ccn2* misexpression (a) and representative H&E images (a’) of left and right proximal tibiae at E17.5 (n=3). **b**, **b’**, Representative images (b, boxed regions shown magnified in I-IV) and quantification (b’) of Ki67 staining in the RZ of CTL and EXP proximal tibia (n=5 and 3, respectively.)

**Extended Data Figure 10.** Comparison of cell proportions between control (CTP) and *Cart-p21* (ETP) samples within the Gli1-lineage (tdTom^+^). GoR, groove of Ranvier. Note how the RestingChon 2 population (arrow) is mainly expanded at the expense of the GoR one.

## Notes

### Summary of Updates

New functional and mechanistic results. Updated narrative. Improved bioinformatics analysis.

## References

1 Wong, M. & Gilmour, D. Getting back on track: exploiting canalization to uncover the mechanisms of developmental robustness. Curr Opin Genet Dev 63, 53–60 (2020). 10.1016/j.gde.2020.04.001

2 Rolian, C. Endochondral ossification and the evolution of limb proportions. Wiley interdisciplinary reviews. Developmental biology, e373 (2020). 10.1002/wdev.373

3 Kronenberg, H. M. Developmental regulation of the growth plate. Nature 423, 332–336 (2003). 10.1038/nature01657

4 Zhou, X. et al. Chondrocytes transdifferentiate into osteoblasts in endochondral bone during development, postnatal growth and fracture healing in mice. PLoS Genet 10, e1004820 (2014). 10.1371/journal.pgen.1004820

5 Maes, C. et al. Osteoblast precursors, but not mature osteoblasts, move into developing and fractured bones along with invading blood vessels. Dev Cell 19, 329–344 (2010). 10.1016/j.devcel.2010.07.010

6 Karlsson, C., Thornemo, M., Henriksson, H. B. & Lindahl, A. Identification of a stem cell niche in the zone of Ranvier within the knee joint. J Anat 215, 355–363 (2009). 10.1111/j.1469-7580.2009.01115.x

7 Newton, P. T. et al. A radical switch in clonality reveals a stem cell niche in the epiphyseal growth plate. Nature 567, 234–238 (2019). 10.1038/s41586-019-0989-6

8 Harper, J. W., Adami, G. R., Wei, N., Keyomarsi, K. & Elledge, S. J. The p21 Cdk-interacting protein Cip1 is a potent inhibitor of G1 cyclin-dependent kinases. Cell 75, 805–816 (1993).

9 Shiratori, H., Yashiro, K., Shen, M. M. & Hamada, H. Conserved regulation and role of Pitx2 in situs-specific morphogenesis of visceral organs. Development 133, 3015–3025 (2006). 10.1242/dev.02470

10 Posey, K. L. et al. An inducible cartilage oligomeric matrix protein mouse model recapitulates human pseudoachondroplasia phenotype. Am J Pathol 175, 1555–1563 (2009). 10.2353/ajpath.2009.090184

11 Rosello-Diez, A., Madisen, L., Bastide, S., Zeng, H. & Joyner, A. L. Cell-nonautonomous local and systemic responses to cell arrest enable long-bone catch-up growth in developing mice. PLoS Biol 16, e2005086 (2018). 10.1371/journal.pbio.2005086

12 Aibar, S. et al. SCENIC: single-cell regulatory network inference and clustering. Nature methods 14, 1083–1086 (2017). 10.1038/nmeth.4463

13 Shi, Y. et al. Gli1 identifies osteogenic progenitors for bone formation and fracture repair. Nature communications 8, 2043 (2017). 10.1038/s41467-017-02171-2

14 Phipson, B. et al. propeller: testing for differences in cell type proportions in single cell data. Bioinformatics 38, 4720–4726 (2022). 10.1093/bioinformatics/btac582

15 Ahmadzadeh, E. et al. A collection of genetic mouse lines and related tools for inducible and reversible intersectional misexpression. Development 147 (2020). 10.1242/dev.186650

16 Ahn, S. & Joyner, A. L. Dynamic changes in the response of cells to positive hedgehog signaling during mouse limb patterning. Cell 118, 505–516 (2004). 10.1016/j.cell.2004.07.023

17 Madisen, L. et al. A robust and high-throughput Cre reporting and characterization system for the whole mouse brain. Nat Neurosci 13, 133–140 (2010). 10.1038/nn.2467

18 Abad, V. et al. The role of the resting zone in growth plate chondrogenesis. Endocrinology 143, 1851–1857 (2002). 10.1210/endo.143.5.8776

19 H’ng, C. H. et al. Compensatory growth and recovery of cartilage cytoarchitecture after transient cell death in fetal mouse limbs. Nature communications 15, 2940 (2024). 10.1038/s41467-024-47311-7

20 Li, B. et al. Gli1 labels progenitors during chondrogenesis in postnatal mice. EMBO reports (2024). 10.1038/s44319-024-00093-x

21 He, M. et al. Strategies and Tools for Combinatorial Targeting of GABAergic Neurons in Mouse Cerebral Cortex. Neuron 92, 555 (2016). 10.1016/j.neuron.2016.10.009

22 Fenichel, I., Evron, Z. & Nevo, Z. The perichondrial ring as a reservoir for precartilaginous cells. In vivo model in young chicks’ epiphysis. Int Orthop 30, 353–356 (2006). 10.1007/s00264-006-0082-2

23 Chung, M. I., Bujnis, M., Barkauskas, C. E., Kobayashi, Y. & Hogan, B. L. M. Niche-mediated BMP/SMAD signaling regulates lung alveolar stem cell proliferation and differentiation. Development 145 (2018). 10.1242/dev.163014

24 Squair, J. W. et al. Confronting false discoveries in single-cell differential expression. Nature communications 12, 5692 (2021). 10.1038/s41467-021-25960-2

25 Guo, F., Carter, D. E. & Leask, A. Mechanical tension increases CCN2/CTGF expression and proliferation in gingival fibroblasts via a TGFbeta-dependent mechanism. PLoS One 6, e19756 (2011). 10.1371/journal.pone.0019756

26 Kubota, S. & Takigawa, M. The role of CCN2 in cartilage and bone development. J. Cell Commun. Signal. 5, 209–217 (2011). 10.1007/s12079-011-0123-5

27 Nishida, T., Maeda, A., Kubota, S. & Takigawa, M. Role of mechanical-stress inducible protein Hcs24/CTGF/CCN2 in cartilage growth and regeneration: mechanical stress induces expression of Hcs24/CTGF/CCN2 in a human chondrocytic cell line HCS-2/8, rabbit costal chondrocytes and meniscus tissue cells. Biorheology 45, 289–299 (2008).

28 Browaeys, R., et al. MultiNicheNet: a flexible framework for differential cell-cell communication analysis from multi-sample multi-condition single-cell transcriptomics data. *bioRxiv* (2023).

29 Takigawa, M. CCN2: a master regulator of the genesis of bone and cartilage. Journal of cell communication and signaling 7, 191–201 (2013). 10.1007/s12079-013-0204-8

30 Mizuhashi, K. et al. Resting zone of the growth plate houses a unique class of skeletal stem cells. Nature 563, 254–258 (2018). 10.1038/s41586-018-0662-5

31 Muruganandan, S. et al. A FoxA2+ long-term stem cell population is necessary for growth plate cartilage regeneration after injury. Nature communications 13, 2515 (2022). 10.1038/s41467-022-30247-1

32 Chan, L. L., Huang, J., Hagiwara, Y., Aguila, L. & Rowe, D. Discriminating multiplexed GFP reporters in primary articular chondrocyte cultures using image cytometry. J Fluoresc 24, 1041–1053 (2014). 10.1007/s10895-014-1383-2

33 Ahn, S. & Joyner, A. L. In vivo analysis of quiescent adult neural stem cells responding to Sonic hedgehog. Nature 437, 894–897 (2005). 10.1038/nature03994

34 Hamilton, T. G., Klinghoffer, R. A., Corrin, P. D. & Soriano, P. Evolutionary divergence of platelet-derived growth factor alpha receptor signaling mechanisms. Mol Cell Biol 23, 4013–4025 (2003). 10.1128/MCB.23.11.4013-4025.2003

35 Pei, W. et al. Polylox barcoding reveals haematopoietic stem cell fates realized in vivo. Nature 548, 456–460 (2017). 10.1038/nature23653

36 Zhang, B. et al. A human embryonic limb cell atlas resolved in space and time. Nature (2023). 10.1038/s41586-023-06806-x

37 McGinnis, C. S., Murrow, L. M. & Gartner, Z. J. DoubletFinder: doublet detection in single- cell RNA sequencing data using artificial nearest neighbors. Cell systems 8, 329–337. e324 (2019).

38 Browaeys, R. et al. MultiNicheNet: a flexible framework for differential cell-cell communication analysis from multi-sample multi-condition single-cell transcriptomics data. BioRxiv, 2023.2006. 2013.544751 (2023).

39 Aibar, S. et al. SCENIC: single-cell regulatory network inference and clustering. Nature methods 14, 1083–1086 (2017).

40 Van de Sande, B. et al. A scalable SCENIC workflow for single-cell gene regulatory network analysis. Nature protocols 15, 2247–2276 (2020).

41 Beltran Diaz, S., et al. A new pipeline to automatically segment and semi-automatically measure bone length on 3D models obtained by Computed Tomography. Frontiers in Cell and Developmental Biology (2021). 10.3389/fcell.2021.736574

