## Supplementary material for "Long-lived chondroprogenitors are generated by Gli1^+^ fetal-limb cells and are replenished upon cell-cycle arrest in the cartilage": Ext. Data Fig.

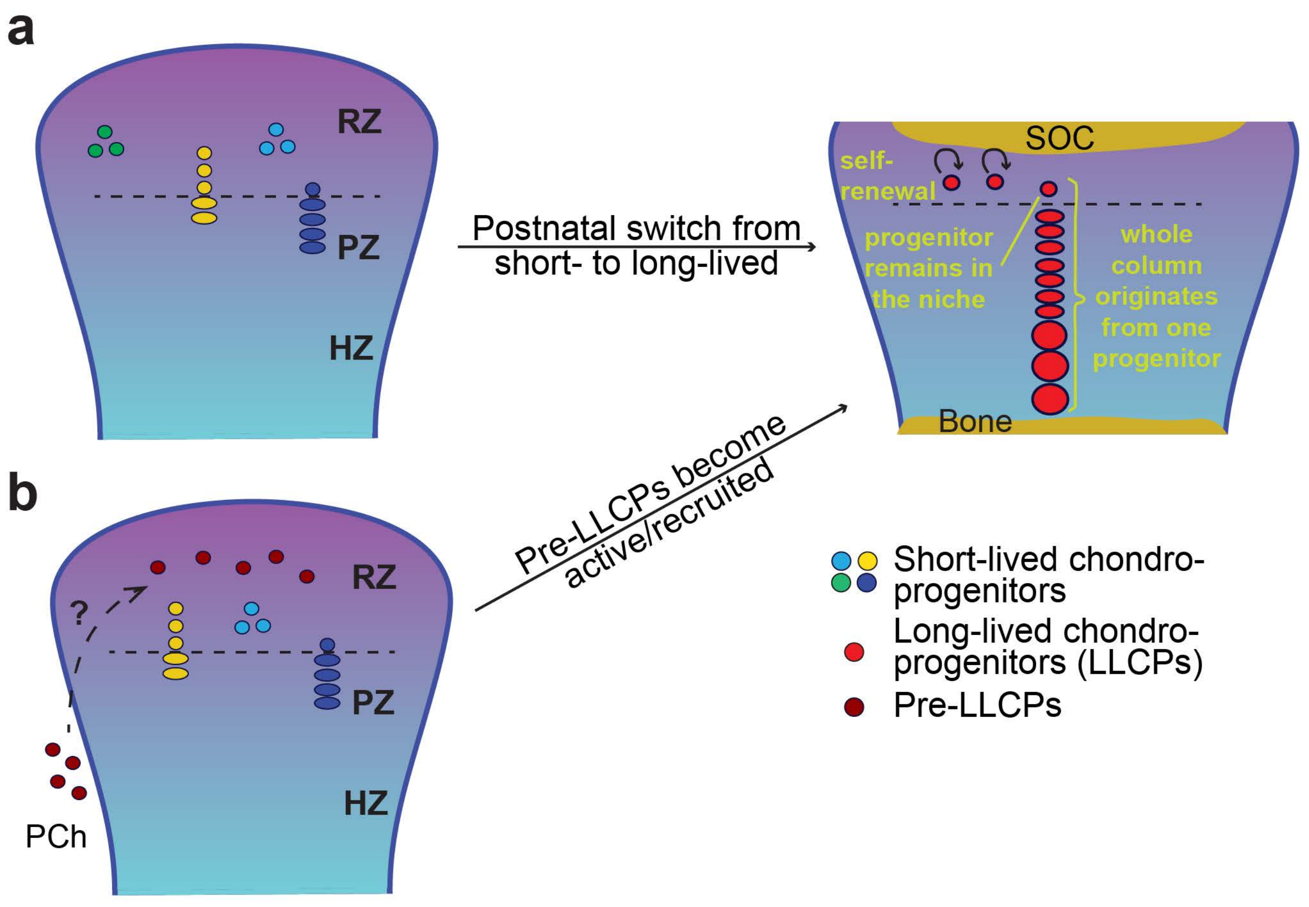
a
RZ
PZ
HZ
Postnatal switch from short- to long-lived
SOC
self-renewal
progenitor remains in the niche
whole column originates from one progenitor
Bone
b
RZ
?
PZ
HZ
PCh
Pre-LLCPs become active/recruited
Short-lived chondro-progenitors
Long-lived chondro-progenitors (LLCPs)
Pre-LLCPs

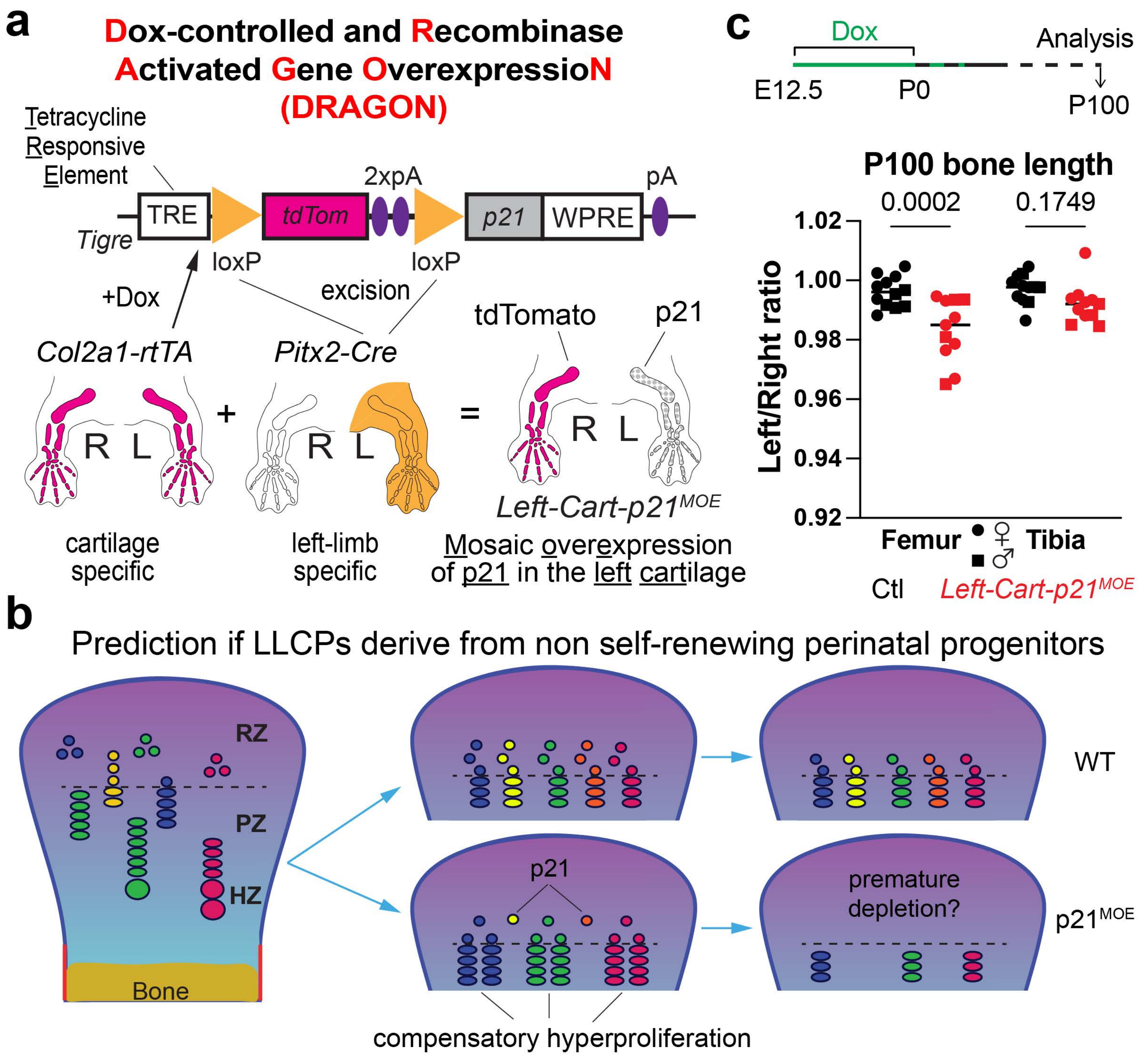
a
Dox-controlled and Recombinase Activated Gene OverexpressioN (DRAGON)
Tetracycline Responsive Element
TRE
tdTom
2xpA
p21
WPRE
pA
Tigre
loxP
loxP
excision
+Dox
Col2a1-rtTA
Pitx2-Cre
tdTomato
p21
R L
R L
R L
Left-Cart-p21^MOE
cartilage specific
left-limb specific
Mosaic overexpression of p21 in the left cartilage
c
Dox
Analysis
E12.5
P0
P100
P100 bone length
0.0002
0.1749
Left/Right ratio
1.02
1.00
0.98
0.96
0.94
0.92
Femur
Tibia
♀
♂
Ctl
Left-Cart-p21^MOE
b
Prediction if LLCPs derive from non self-renewing perinatal progenitors
RZ
PZ
HZ
Bone
WT
p21
premature depletion?
p21^MOE
compensatory hyperproliferation

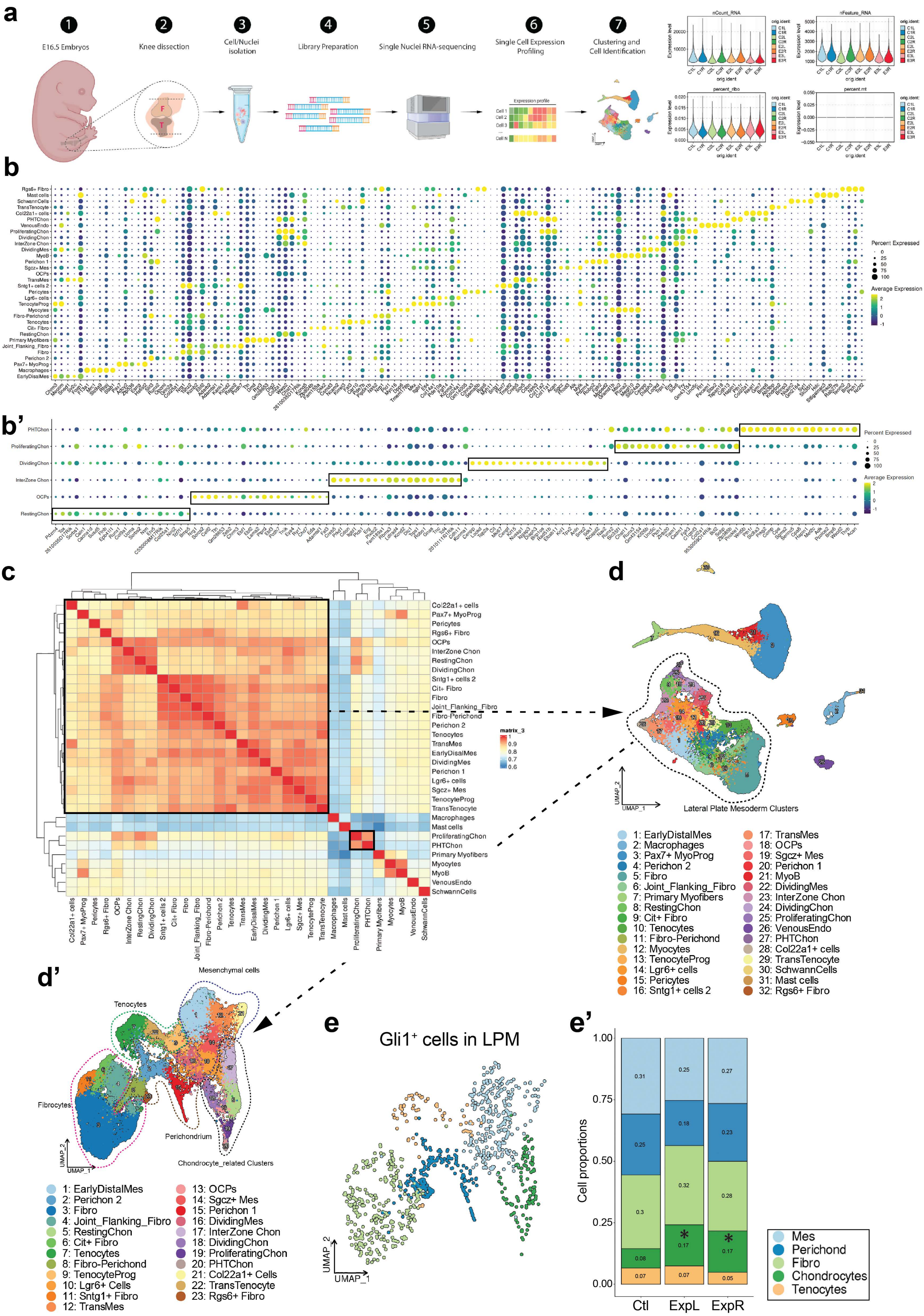

a
E16.5 Embryos
Knee dissection
Cell/Nuclei isolation
Library Preparation
Single Nuclei RNA-sequencing
Single Cell Expression Profiling
Clustering and Cell Identification
Expression profile
Cell 1
Cell 2
Cell 3
Cell N
nCount_RNA
nFeature_RNA
percent_ribo
percent.mt
Expression level
orig.ident
C1L C1R C2L C2R E2L E2R E3L E3R
b
Percent Expressed
Average Expression
b'
PHTChon
ProliferatingChon
DividingChon
InterZone Chon
OCPs
RestingChon
c
matrix_3
d
Lateral Plate Mesoderm Clusters
UMAP_1
UMAP_2
1: EarlyDistalMes
2: Macrophages
3: Pax7+ MyoProg
4: Perichon 2
5: Fibro
6: Joint_Flanking_Fibro
7: Primary Myofibers
8: RestingChon
9: Cit+ Fibro
10: Tenocytes
11: Fibro-Perichond
12: Myocytes
13: TenocyteProg
14: Lgr6+ cells
15: Pericytes
16: Sntg1+ cells 2
17: TransMes
18: OCPs
19: Sgcz+ Mes
20: Perichon 1
21: MyoB
22: DividingMes
23: InterZone Chon
24: DividingChon
25: ProliferatingChon
26: VenousEndo
27: PHTChon
28: Col22a1+ cells
29: TransTenocyte
30: SchwannCells
31: Mast cells
32: Rgs6+ Fibro
d'
Mesenchymal cells
Tenocytes
Fibrocytes
Perichondrium
Chondrocyte_related Clusters
1: EarlyDistalMes
2: Perichon 2
3: Fibro
4: Joint_Flanking_Fibro
5: RestingChon
6: Cit+ Fibro
7: Tenocytes
8: Fibro-Perichond
9: TenocyteProg
10: Lgr6+ Cells
11: Sntg1+ Fibro
12: TransMes
13: OCPs
14: Sgcz+ Mes
15: Perichon 1
16: DividingMes
17: InterZone Chon
18: DividingChon
19: ProliferatingChon
20: PHTChon
21: Col22a1+ Cells
22: TransTenocyte
23: Rgs6+ Fibro
e
Gli1⁺ cells in LPM
e'
Cell proportions
Ctl
ExpL
ExpR
Mes
Perichon
Fibro
Chondrocytes
Tenocytes

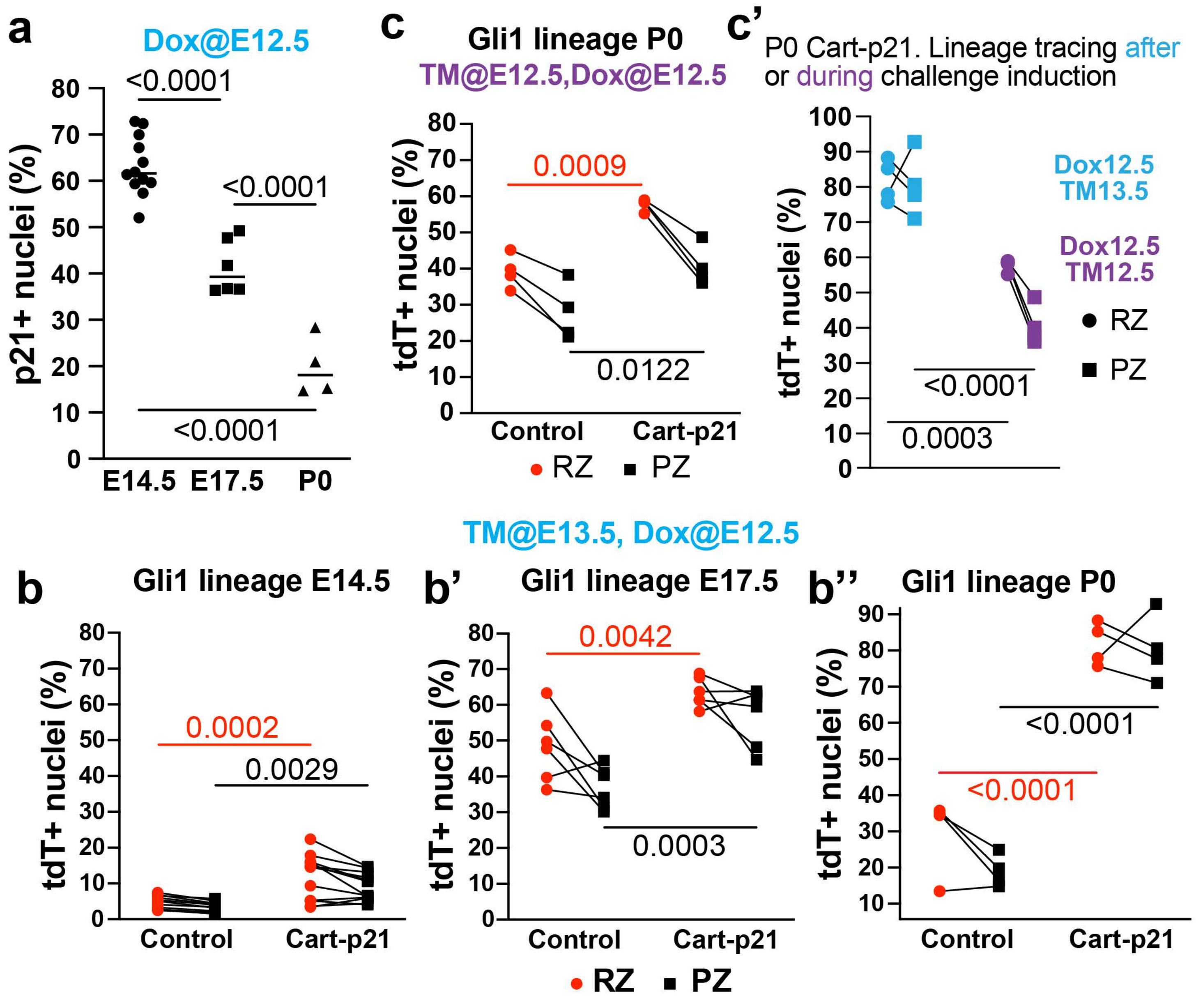
a
Dox@E12.5
p21+ nuclei (%)
<0.0001
<0.0001
<0.0001
E14.5
E17.5
P0
c
Gli1 lineage P0
TM@E12.5,Dox@E12.5
tdT+ nuclei (%)
0.0009
0.0122
Control
Cart-p21
RZ
PZ
c'
P0 Cart-p21. Lineage tracing after or during challenge induction
tdT+ nuclei (%)
Dox12.5 TM13.5
Dox12.5 TM12.5
RZ
PZ
<0.0001
0.0003
TM@E13.5, Dox@E12.5
b
Gli1 lineage E14.5
tdT+ nuclei (%)
0.0002
0.0029
Control
Cart-p21
b'
Gli1 lineage E17.5
tdT+ nuclei (%)
0.0042
0.0003
Control
Cart-p21
b''
Gli1 lineage P0
tdT+ nuclei (%)
<0.0001
<0.0001
Control
Cart-p21
RZ
PZ

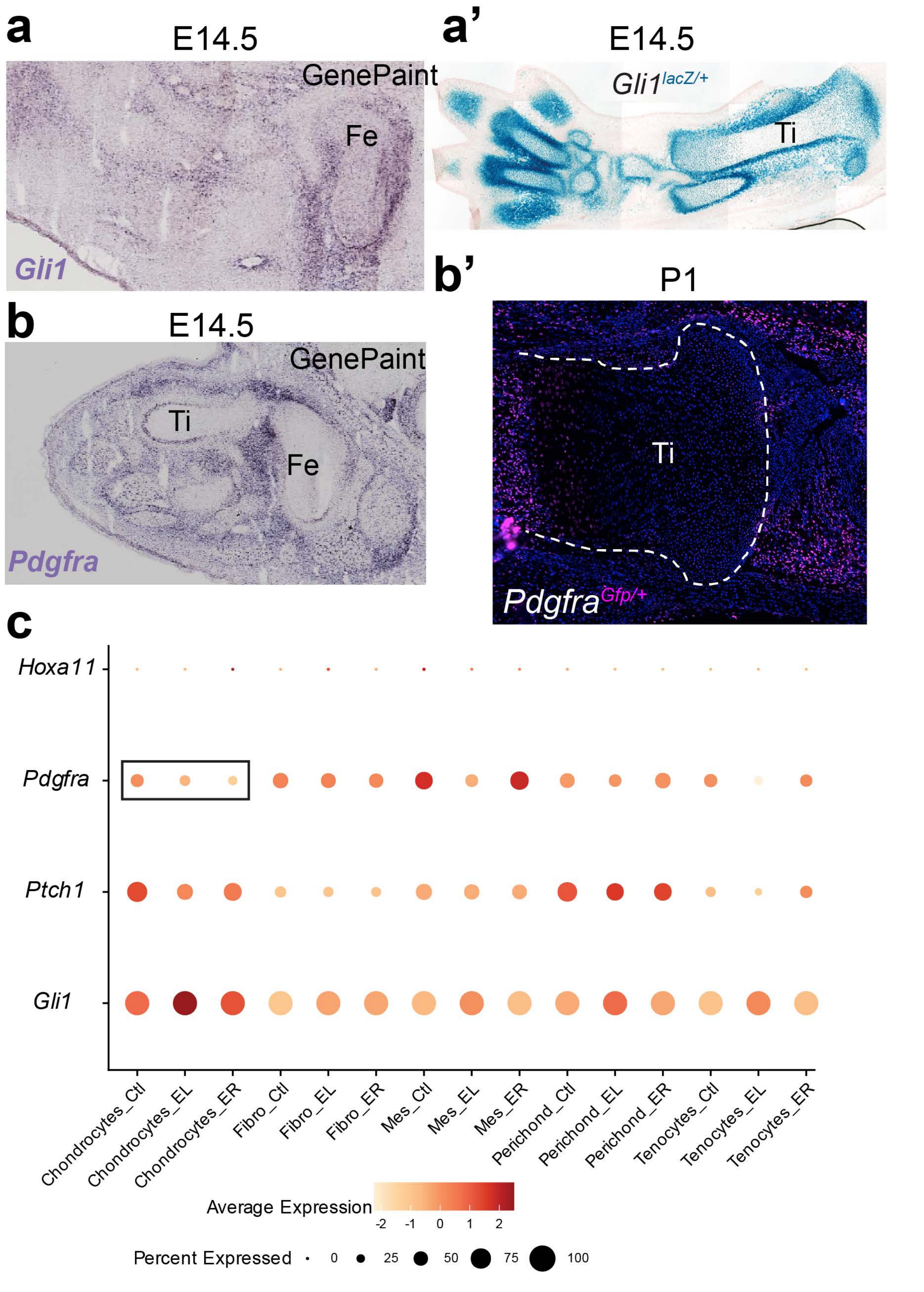
a
E14.5
GenePaint
Fe
Gli1
a'
E14.5
Gli1^lacZ/+
Ti
b
E14.5
GenePaint
Ti
Fe
Pdgfra
b'
P1
Ti
Pdgfra^Gfp/+
c
Hoxa11
Pdgfra
Ptch1
Gli1
Chondrocytes_Ctl
Chondrocytes_EL
Chondrocytes_ER
Fibro_Ctl
Fibro_EL
Fibro_ER
Mes_Ctl
Mes_EL
Mes_ER
Perichond_Ctl
Perichond_EL
Perichond_ER
Tenocytes_Ctl
Tenocytes_EL
Tenocytes_ER
Average Expression
-2 -1 0 1 2
Percent Expressed
0 25 50 75 100

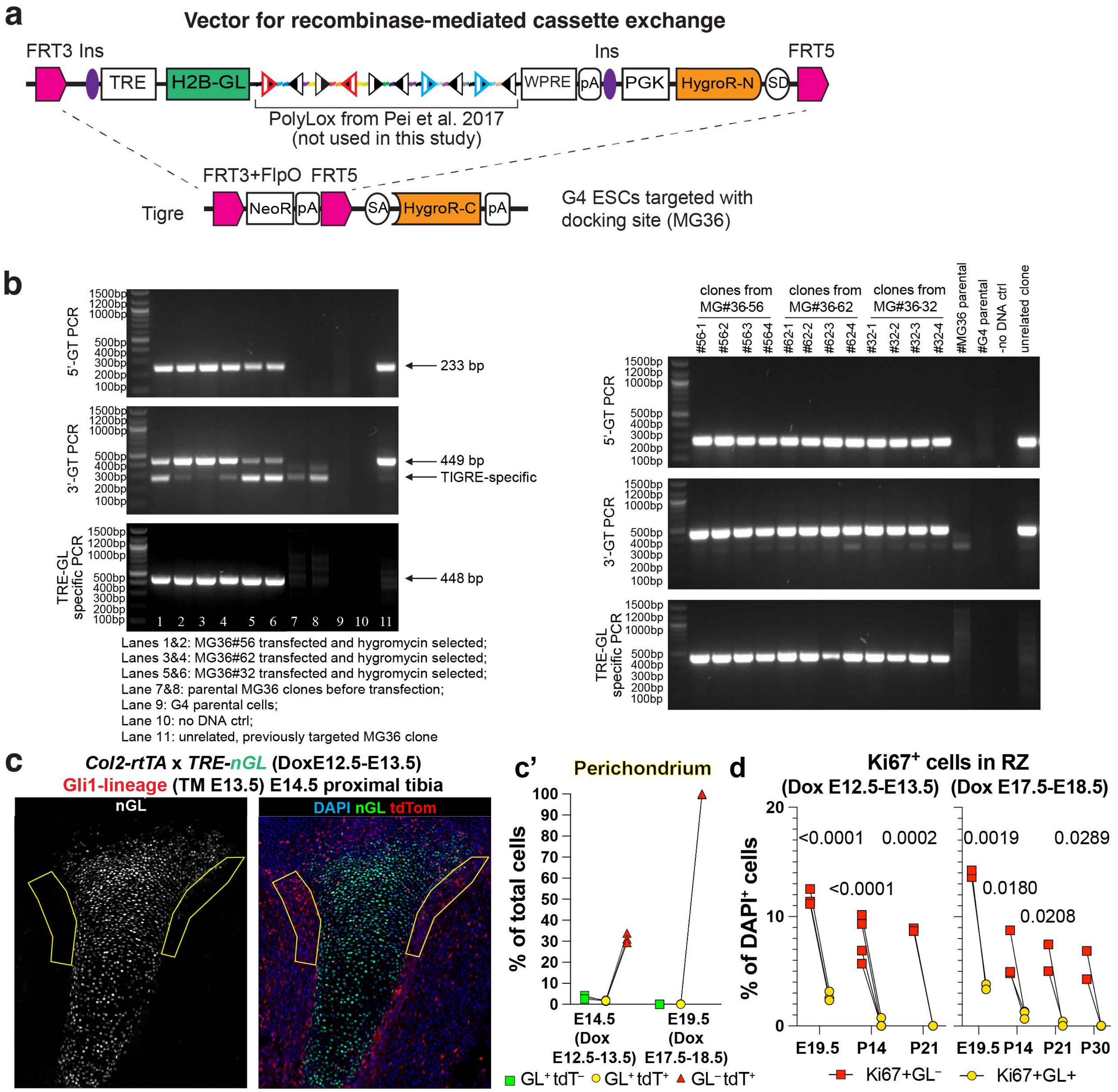
a
Vector for recombinase-mediated cassette exchange
FRT3 Ins
TRE
H2B-GL
WPRE
pA
Ins
PGK
HygroR-N
SD
FRT5
PolyLox from Pei et al. 2017
(not used in this study)
Tigre
FRT3+FlpO
NeoR
pA
FRT5
SA
HygroR-C
pA
G4 ESCs targeted with
docking site (MG36)
b
5'-GT PCR
3'-GT PCR
TRE-GL specific PCR
1500bp
1200bp
1000bp
500bp
400bp
300bp
200bp
100bp
233 bp
449 bp
TIGRE-specific
448 bp
1 2 3 4 5 6 7 8 9 10 11
Lanes 1&2: MG36#56 transfected and hygromycin selected;
Lanes 3&4: MG36#62 transfected and hygromycin selected;
Lanes 5&6: MG36#32 transfected and hygromycin selected;
Lane 7&8: parental MG36 clones before transfection;
Lane 9: G4 parental cells;
Lane 10: no DNA ctrl;
Lane 11: unrelated, previously targeted MG36 clone
clones from MG#36-56
clones from MG#36-62
clones from MG#36-32
#56-1
#56-2
#56-3
#56-4
#62-1
#62-2
#62-3
#62-4
#32-1
#32-2
#32-3
#32-4
#MG36 parental
#G4 parental
-no DNA ctrl
unrelated clone
c
Col2-rtTA x TRE-nGL (DoxE12.5-E13.5)
Gli1-lineage (TM E13.5) E14.5 proximal tibia
nGL
DAPI nGL tdTom
c'
Perichondrium
% of total cells
100
90
80
70
60
50
40
30
20
10
0
E14.5 (Dox E12.5-13.5)
E19.5 (Dox E17.5-18.5)
GL+ tdT−
GL+ tdT+
GL− tdT+
d
Ki67+ cells in RZ
(Dox E12.5-E13.5)
(Dox E17.5-E18.5)
% of DAPI+ cells
20
10
0
<0.0001
0.0002
<0.0001
0.0019
0.0289
0.0180
0.0208
E19.5
P14
P21
P30
Ki67+GL−
Ki67+GL+

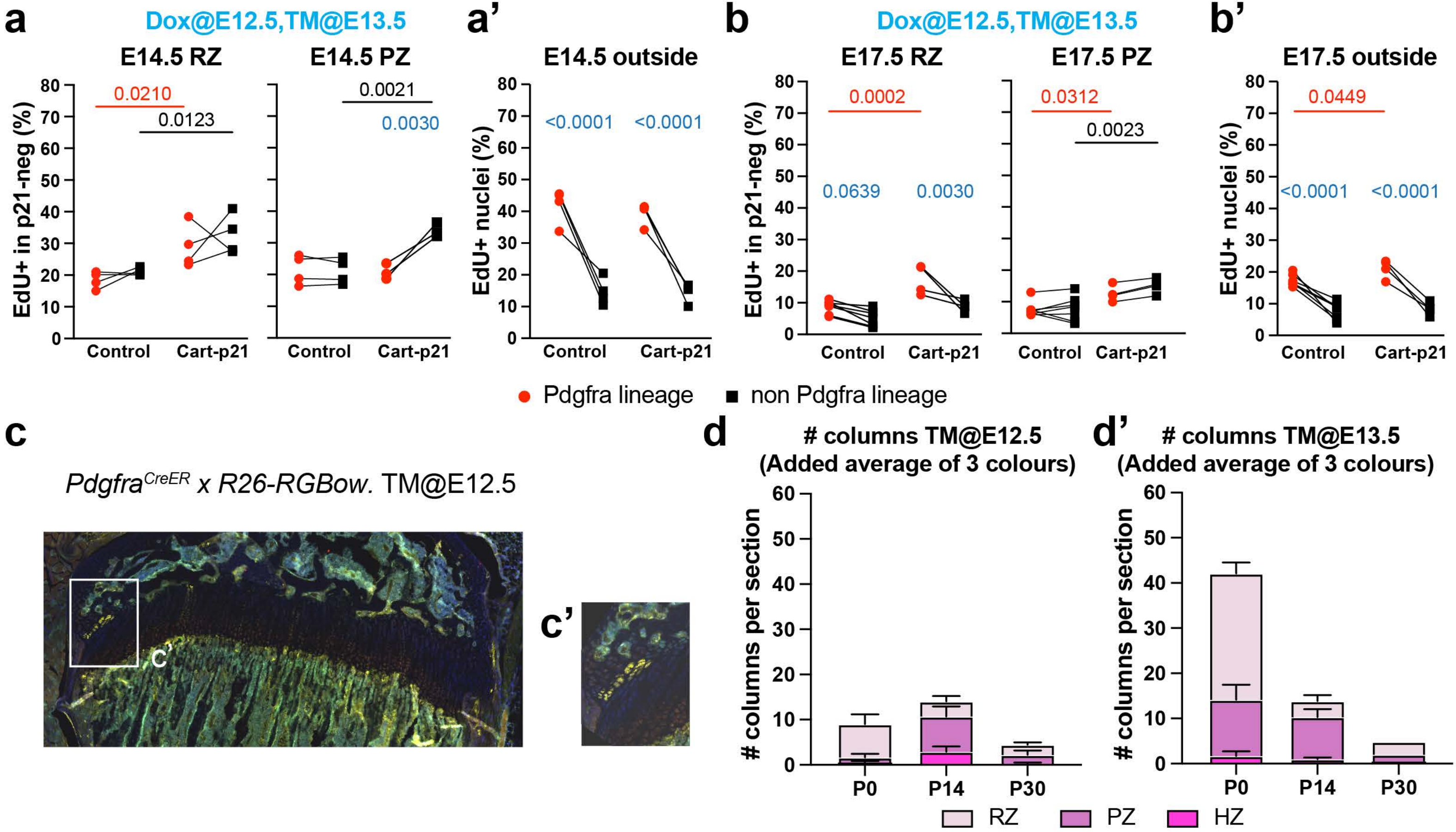

a
Dox@E12.5,TM@E13.5
E14.5 RZ
E14.5 PZ
EdU+ in p21-neg (%)
0.0210
0.0123
0.0021
0.0030
Control
Cart-p21
a'
E14.5 outside
EdU+ nuclei (%)
<0.0001
<0.0001
b
Dox@E12.5,TM@E13.5
E17.5 RZ
E17.5 PZ
0.0002
0.0639
0.0030
0.0312
0.0023
b'
E17.5 outside
0.0449
<0.0001
<0.0001
Pdgfra lineage
non Pdgfra lineage
c
Pdgfra^CreER x R26-RGBow. TM@E12.5
C'
c'
d
### columns TM@E12.5
(Added average of 3 colours)
d'
### columns TM@E13.5
(Added average of 3 colours)
### columns per section
P0
P14
P30
RZ
PZ
HZ

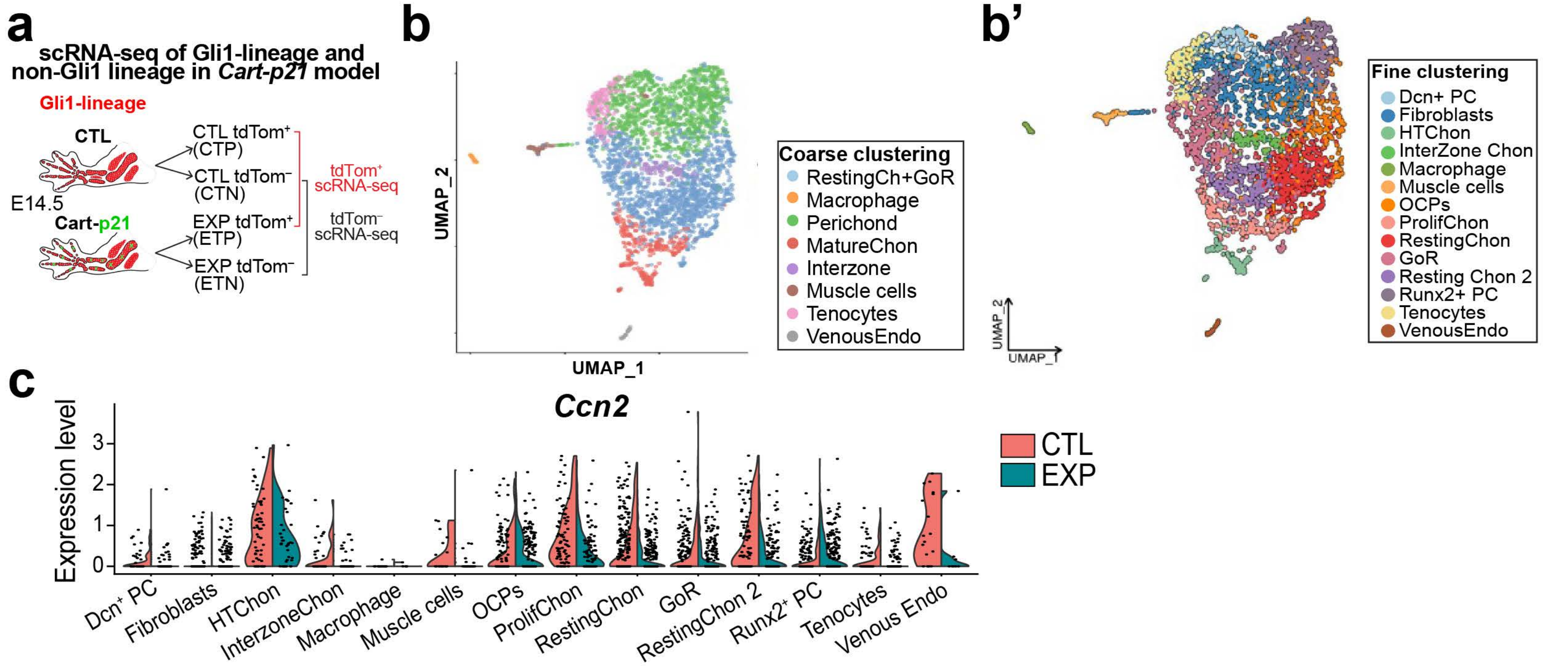


**d** GO terms related to *Ccn2* (EXP vs. CTL)

| Pathway | NES | pval | padj |
|---|---|---|---|
| GOBP_CELLULAR_RESPONSE_TO_STRESS | -1.31 | 7.8e-04 | 3.1e-02 |
| GOBP_CELL_CYCLE | -1.46 | 2.5e-04 | 2.2e-02 |
| GOBP_CELL_CYCLE_PROCESS | -1.52 | 2.5e-04 | 2.2e-02 |
| GOBP_REGULATION_OF_CELL_CYCLE | -1.55 | 2.4e-04 | 2.2e-02 |
| GOBP_POSITIVE_REGULATION_OF_CELL_CYCLE | -1.56 | 6.8e-04 | 3.0e-02 |
| GOBP_SKELETAL_SYSTEM_MORPHOGENESIS | -1.63 | 8.9e-04 | 3.4e-02 |
| GOBP_REGULATION_OF_CELL_CYCLE_PROCESS | -1.67 | 2.3e-04 | 2.2e-02 |
| GOBP_SKELETAL_SYSTEM_DEVELOPMENT | -1.67 | 2.3e-04 | 2.2e-02 |
| GOBP_DNA_METABOLIC_PROCESS | -1.68 | 2.4e-04 | 2.2e-02 |
| GOCC_EXTERNAL_ENCAPSULATING_STRUCTURE | -1.70 | 2.2e-04 | 2.2e-02 |
| GOBP_EXTRACELLULAR_STRUCTURE_ORGANIZATION | -1.75 | 2.3e-04 | 2.2e-02 |
| GOBP_CONNECTIVE_TISSUE_DEVELOPMENT | -1.93 | 2.3e-04 | 2.2e-02 |
| GOBP_CHONDROCYTE_DIFFERENTIATION | -1.96 | 2.2e-04 | 2.2e-02 |
| GOBP_CARTILAGE_DEVELOPMENT | -2.11 | 2.2e-04 | 2.2e-02 |

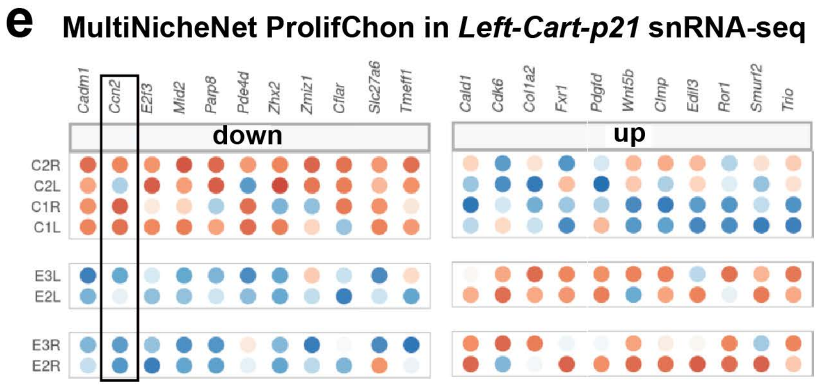


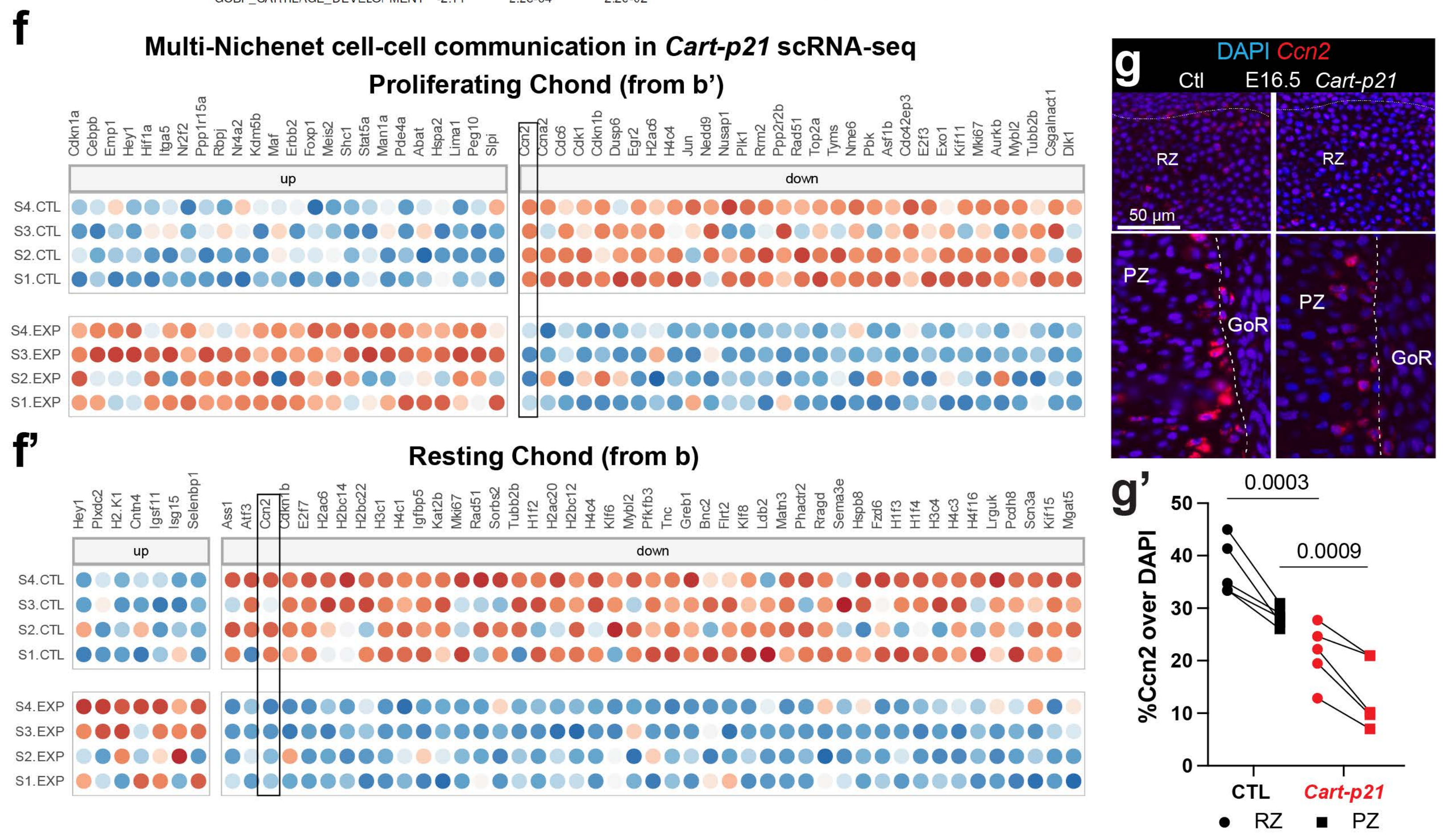

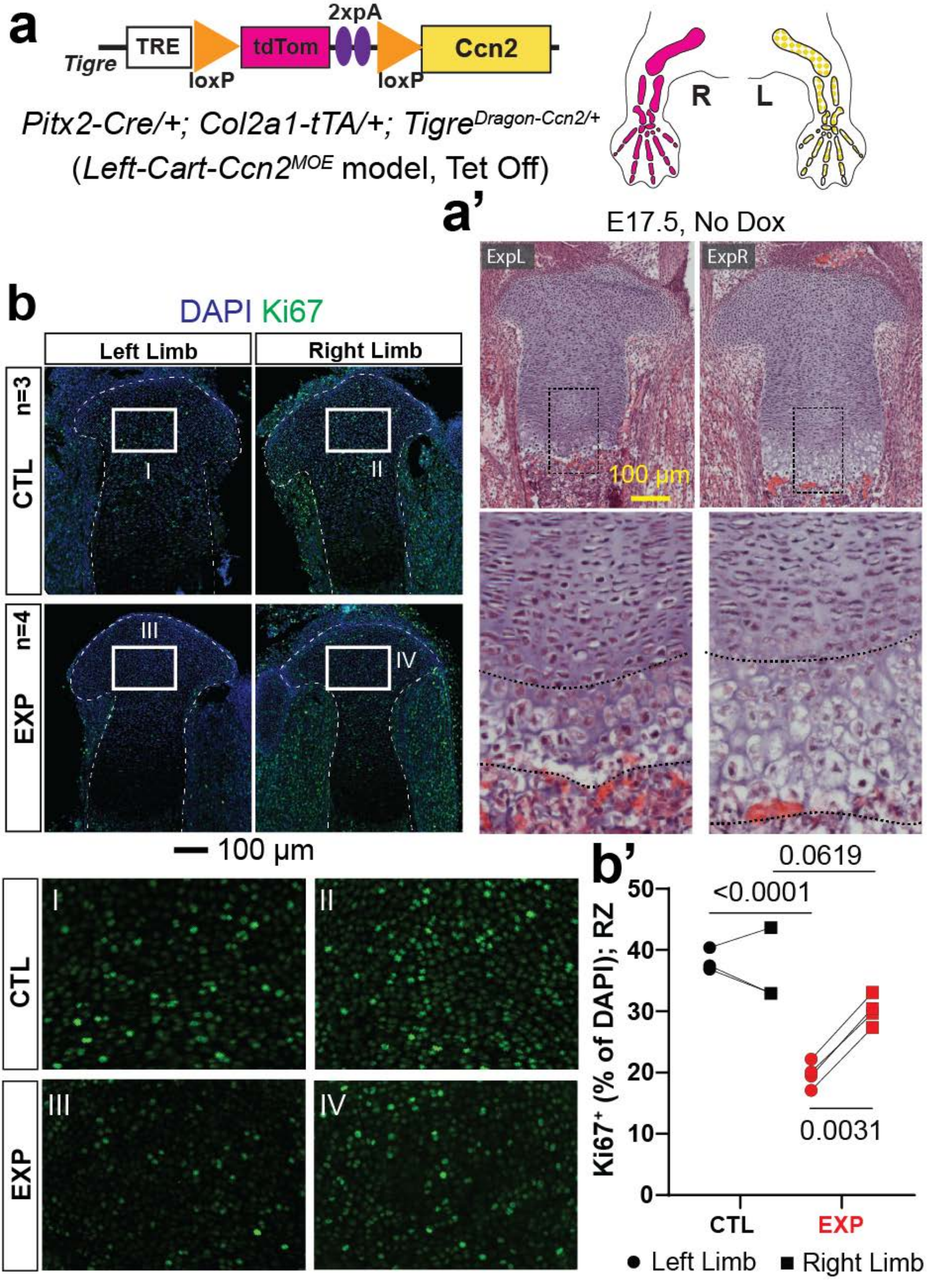
a
Tigre
TRE
loxP
tdTom
2xpA
loxP
Ccn2
Pitx2-Cre/+; Col2a1-tTA/+; TigreDragon-Ccn2/+
(Left-Cart-Ccn2MOE model, Tet Off)
R
L
a'
E17.5, No Dox
ExpL
ExpR
100 µm
b
DAPI Ki67
Left Limb
Right Limb
n=3
CTL
I
II
n=4
EXP
III
IV
100 µm
CTL
EXP
I
II
III
IV
b'
Ki67+ (% of DAPI); RZ
0.0619
<0.0001
0.0031
CTL
EXP
Left Limb
Right Limb

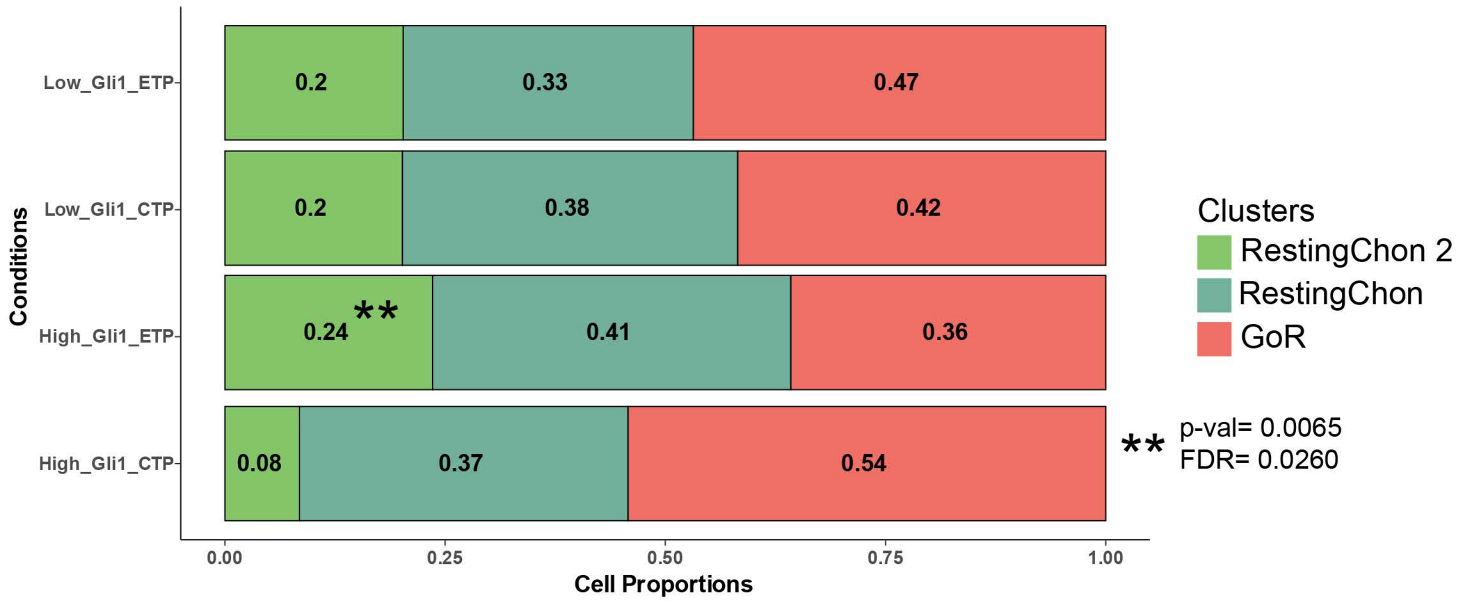

Low_Gli1_ETP
Low_Gli1_CTP
High_Gli1_ETP
High_Gli1_CTP
Conditions
0.2
0.33
0.47
0.2
0.38
0.42
0.24
**
0.41
0.36
0.08
0.37
0.54
0.00
0.25
0.50
0.75
1.00
Cell Proportions
Clusters
RestingChon 2
RestingChon
GoR
**
p-val= 0.0065
FDR= 0.0260
